# Mutational scanning reveals substrate-assisted autoregulation of the WNT destruction complex

**DOI:** 10.1101/2025.10.17.683169

**Authors:** Murugesh Padmanarayana, Saira Sakalas, Parijat Sarkar, Mengxiao Ma, Ethan R. Garvin, Ethan Lee, Steven M. Corsello, Sebastian Guettler, Ganesh V. Pusapati, Rajat Rohatgi

## Abstract

The β-catenin destruction complex (BDC) is a central node in WNT/β-catenin signaling, governing embryonic development and adult tissue homeostasis. Although recognized as a prime therapeutic target in colorectal cancer (CRC) for three decades, its dynamic architecture and biochemical complexity have hindered mechanistic understanding. Here, we systematically mapped the sequence-function landscape of the BDC using tiled base editor screens across four endogenous components—*CTNNB1*, *AXIN1*, *APC*, and *GSK3B*. Validation studies identified ∼150 previously unreported mutations across these genes that affected WNT/β-catenin signaling. In addition to known cancer-associated mutations, we discovered rare gain-of-function and separation-of-function alleles of *AXIN1* and *CTNNB1* that provide mechanistic insights into complex assembly and regulation. We describe a region in β-catenin that regulates its binding to TCF/LEF transcription factors and demonstrate that the AXIN1–β-catenin interface is critical for controlling signaling flux through the oncogenic BDC. Mechanistic studies revealed that assembly of the oncogenic BDC is scaffolded by its own substrate β-catenin, establishing an autoregulatory mechanism that represents an unexploited vulnerability in cancers harboring common APC truncations. Our comprehensive mutational resource provides a foundation for understanding WNT/β-catenin signaling mechanisms in health and disease, while revealing strategies for therapeutic intervention in WNT-driven cancers.

## Introduction

The WNT/β-catenin pathway is a cell-cell communication system that coordinates the patterning and morphogenesis of most tissues during development^1,2^. Even minor defects in signaling strength cause multiorgan birth defects in humans^3^. In adults, WNT/β-catenin signaling maintains tissue homeostasis by guiding regeneration and repair and has been implicated in chronic degenerative diseases of multiple tissues, including the brain, bone and gastrointestinal system^1^. Unchecked WNT/β-catenin signaling is a major cause of human cancer morbidity and mortality, yet we lack a single FDA-approved agent targeting this pathway in any cancer^4^. Truncating mutations in the tumor suppressor gene Adenomatous Polyposis Coli (*APC*) lead to ligand-independent, oncogenic signaling that drives >80% of human colorectal cancer (CRC), the second leading cause of cancer deaths^5–9^.

APC is a component of the <u>β</u>-catenin <u>d</u>estruction <u>c</u>omplex (BDC), a multi-protein complex that limits WNT/β-catenin signaling by promoting the degradation of β-catenin, a multi-functional protein that coordinates the WNT gene expression program^10–12^. The BDC is composed of two scaffolding proteins (APC and AXIN1), two kinases (CK1α and GSK3β), and β-catenin. In the absence of WNTs, the BDC keeps signaling off by promoting the sequential phosphorylation of β-catenin by CK1α and GSK3β, thus marking it for ubiquitination and degradation^13^ (**Fig.1a**). WNTs promote inactivation of the BDC by the receptor complex^14^. As a result, β-catenin accumulates and assembles an activating enhanceosome complex, including BCL9 and TCF/LEF factors, at target gene promoters^15^.

**Fig. 1.**
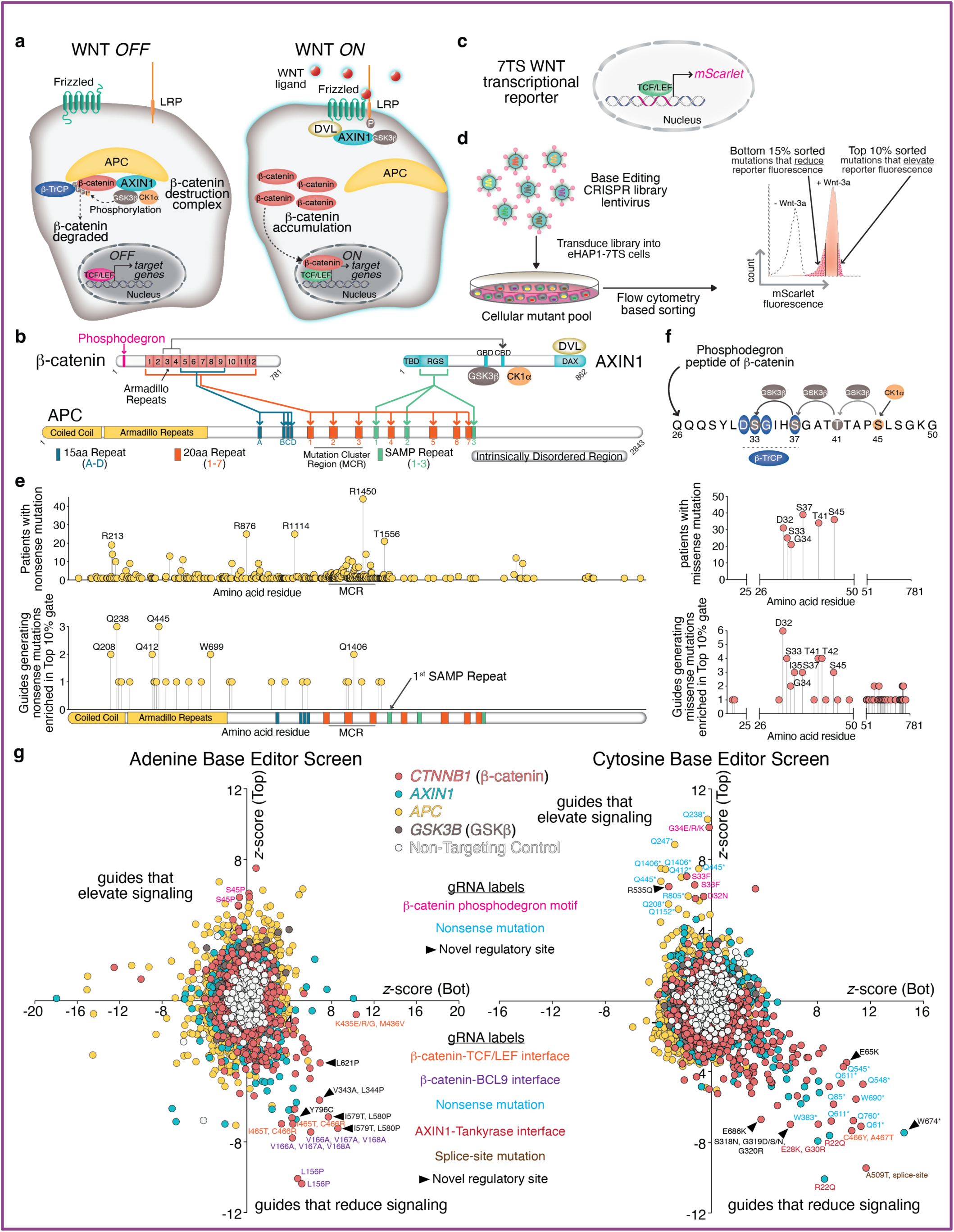
Base editor screens identify mutational hotspots in the β-catenin destruction complex. **a,** Inactivation of the β-catenin destruction complex (BDC) by WNTs leads to the accumulation of β-catenin and activation of WNT target genes. **b,** The BDC (composed of APC, β-catenin, AXIN1 and the AXIN1-bound kinases CK1α and GSK3β) is assembled by multivalent, redundant interactions between folded domains (armadillo repeats, RGS) and linear peptide motifs (15R, 20R, or SAMP). Colored lines indicate previously mapped PPIs between AXIN1-APC, APC-β-catenin, and AXIN1-β-catenin. Most oncogenic mutations in APC truncate the protein in the Mutation Cluster Region (MCR), eliminating all direct contacts with AXIN1 and a subset of contacts with β-catenin. TBD, Tankyrase Binding Domain; DVL, Dishevelled; CBD, β-catenin Binding Domain; GBD, GSK3β Binding Domain, DAX, Domain present in DVL and AXIN. **c,** The transcriptional reporter used to measure WNT/β-catenin signaling strength contains a TCF-responsive element (7 TCF binding sites) driving mScarlet expression (abbreviated 7TS). **d,** Strategy for pooled base-editing screens in human haploid cells carrying the fluorescent WNT reporter (eHAP1-7TS) shown in **c**. **e,** Lollipop plots compare the position (x-axis) and frequency (y-axis) of nonsense mutations in APC from The Cancer Genome Pan-Cancer Atlas (top) with the position of nonsense mutations enriched in the Top 10% gate of our cytosine base editor (CBE) screen (bottom). Most enriched nonsense mutations were found before the first SAMP repeat (see **b**) in both human cancers and our screens. **f,** Lollipop plots show the high frequency of WNT-activating mutations found in β-catenin phosphodegron motif both in human cancers (top) and missense mutations enriched in the Top 10% gate of our base editing screens (bottom). **g,** Plots showing *z*-scores for enrichment of each sgRNA in the Bottom 15% (Bot, x-axis) and Top 10% (Top, y-axis) gates used for sorting mutant cells with diminished or enhanced WNT/β-catenin signaling strength, respectively (see **d**). Each sgRNAs is represented by a single dot colored according to the targeted gene. Label colors identify guides predicted to cause nonsense mutations, splice-site mutations, or mutations at known residues that mediate key PPI or PTMs. Arrowheads indicate sgRNAs that nominate new regulatory sites. See also Extended Data Figures 1, 2, and 3; Supplementary Tables 1 and 3.

Three decades of structural and biochemical studies have mapped multiple, overlapping and seemingly redundant protein-protein interactions (PPIs) between the BDC components^2,10,11,16^ (**Fig.1b**). β-catenin contains a structured central armadillo (ARM) domain composed of 12 repeats and flexible N- and C-terminal segments. β-catenin engages both AXIN1 and APC (and the TCF/LEF family of transcription factors in the nucleus) in a groove that extends along the ARM superhelix. While AXIN1 binds at ARM repeats 3-4, APC and the TCF/LEF proteins bind along a much longer extent of this groove. APC can bind to β-catenin through four 15aa (15R A-D) and six 20aa (20R 1-7, excluding 20R2) peptide motifs. These motifs in APC can bind β-catenin *in vitro* with varying affinities. Phosphorylation of the 20R repeats by CK1α and GSK3β can enhance their affinities for β-catenin by ∼1000-fold^17,18^. APC contains three SAMP motifs that allow it to independently interact with an N-terminal RGS domain in AXIN1^19^. AXIN1 recruits the kinases CK1α and GSK3β into the BDC complex, bringing them into the proximity of their substrates β-catenin and APC^20–22^. The above picture is further complicated by the fact that other proteins occupy the same groove along the β-catenin ARM domain: the TCF/LEF family of transcription factors in the nucleus, cadherins at adherens junction, and ICAT (inhibitor of β-catenin and TCF4) (reviewed in ^11^). Thus, β-catenin participates in at least three cellular assemblies: adherens junction, the BDC in the cytoplasm that regulates its destruction, and the transcriptional complex in the nucleus that shapes the WNT/β-catenin gene expression program.

The complex biochemistry of the BDC has presented obstacles to both mechanistic understanding and therapeutic targeting. For example, we lack information about which sequence elements and protein surfaces are relevant for the activity of the wild-type and oncogenic BDC *in vivo* under endogenous expression levels. Current models are based predominantly on structural or biochemical data with purified protein fragments or overexpression studies in cells (with the notable exception of causal disease variants identified through genetics). The issue of abundance is particularly relevant for the BDC since its size and composition can vary as concentrations of the individual components change^16^. Here we report adenine and cytosine base editor screens to evaluate the WNT signaling phenotype of mutations introduced across endogenous gene loci encoding BDC components^23–27^. Our screens identified classes of mutations that are highly informative (and thus sought after) in sequence-function studies: activating mutations, separation-of-function mutations and mutations that nominate novel regulatory sites. This strategy enabled the identification of surfaces in component proteins that can be used to regulate the activity of the wild-type and oncogenic BDC.

## Results

### Tiled base editor screens targeting endogenous destruction complex genes

We used promiscuous PAM-flexible (NG-PAM Cas9n) adenine and cytosine base editors (NG-ABE8e and NG-BE3) to generate transition mutations across all exons of endogenous *APC, AXIN1, CTNNB1* (encoding β-catenin) and *GSK3B* (encoding GSK3β) in a pooled format^25–28^. Our ABE and CBE libraries are predicted to generate missense mutations at 80-90% of amino acid residues encoded by each gene (**Extended Data Fig.1a, Supplementary Table 1**). Both libraries also generated silent and splice site mutations, since the design strategy allowed single-guide RNAs (sgRNAs) to extend ∼20 nucleotides into introns at each intron-exon junction (**Extended Data Fig.1b**). However, nonsense mutations will only be generated in the CBE library. The sgRNA library covers each exon in a tiled fashion (**Extended Data Fig.1c**).

Mutagenesis was conducted in haploid human cells (eHAP1) stably expressing our fluorescent transcriptional reporter of WNT/β-catenin signaling strength (7TS, **Fig.1c**)^29^. HAP1 cells, previously established as a powerful model for genetic screens targeting the WNT/β-catenin pathway^29^, only possess one allele of each gene and hence simplify the phenotypic evaluation of endogenous mutations^30–32^. Pooled ABE and CBE mutant cell libraries were treated (separately, in duplicate) with the ligand Wnt-3a. Cells with 7TS fluorescence intensities in the top 10% or bottom 15% of the mutant population were isolated using fluorescence activated cell sorting (FACS) to enrich cells carrying mutations that increased or decreased signaling strength, respectively^29^ (**Fig.1d**). We sequenced the top 10% sorted (Top 10%), bottom 15% sorted (Bottom 15%), and unsorted cell populations using the Illumina platform. (**Extended Data Fig.1d, Supplementary Table 1**). Two z-scores for each sgRNA (for enrichment in the Top 10% and Bottom 15% gates) were calculated based on a distribution of 282 non-targeting control sgRNAs (**Supplementary Table 1**). sgRNAs that cause mutations that elevate signaling would be depleted from the Bottom 15% gate (negative z-score) and enriched in the Top 10% gate (positive z-score). Conversely sgRNAs that reduce signaling would be enriched in the Bottom 15% gate but depleted from the Top 10% gate. Full results of the screen, including plots showing the position of statistically significant sgRNAs along the protein sequence of each BDC component are shown in **Supplementary Table 1** and **Extended Data Figs.2 and 3**. Our untargeted screens identified many known functional sites, including the phosphodegron in β-catenin and the tankyrase, β-catenin and Dishevelled (DVL) binding sites in AXIN1 (**Extended Data Figs.2 and 3).**

Some of the most statistically significant sgRNAs in our screen encode mutations that drive oncogenic WNT/β-catenin signaling in human cancer^7,33^. Over 80% of sporadic and familial CRC cases are driven by truncating mutations that map to a ∼250 amino-acid segment in the central Intrinsically Disordered Region (IDR) of APC known as the Mutation Cluster Region or MCR^34–40^ (**Fig.1e)**. Mutant *APC^mcr^* alleles in human cancer produce a truncated protein rather than eliminating APC from the cell. APC^mcr^ lacks all three SAMP repeats that mediate the direct interaction of APC with AXIN1 (**Fig.1b** and **1e**), thus forming a compromised, oncogenic BDC that cannot fully suppress β-catenin abundance and WNT target gene expression in the absence of ligand stimulation^16,41,42^. Multiple CBE sgRNAs predicted to introduce nonsense mutations in APC were enriched in the Top 10% gate, but all these targeted residues are located before the first SAMP repeat (**Fig.1e**). This result mirrors the observation that truncated APC variants that include the first SAMP are not tumorigenic in mouse models^43^.

The most common oncogenic β-catenin mutations across multiple human cancers disrupt an N-terminal degron motif (^32^DSGIHSGATTAPS^45^) sequentially phosphorylated by two AXIN1-bound kinases, Casein Kinase 1α (CK1α) and GSK-3β^13^ (**Fig.1f**). GSK-3β phosphorylation of Ser33 and Ser37 in this degron generates a high-affinity binding site (^32^DpSGφXpS^37^) for the SCF^β-TrCP^ E3 ligase, which ubiquitylates β-catenin and earmarks it for proteasomal degradation^44^. Multiple overlapping sgRNAs targeting nearly all the critical residues in this phosphodegron and β-TrCP binding site (e.g. the invariant Asp32) were enriched in the Top 10% sorted population, consistent with their expected effect of causing mutations that drive constitutive, high-level WNT/β-catenin signaling (**Fig.1f**).

An *x-y* plot of sgRNA z-scores for enrichment in the Bottom 15% gate (*x-*axis) vs. Top 10% gate (*y*-axis) identified sgRNAs predicted to make mutations in regions known to be critical for BDC function and regulation (**Fig.1g**). AXIN1 mutations in residues that mediate its interaction with the negative regulator tankyrase are predicted to impair WNT/β-catenin signaling^45^. Indeed, sgRNAs encoding such mutations at the AXIN1-tankyrase interface are depleted from the Top 10% gate and enriched in the Bottom 15% gate. sgRNAs predicted to mutate β-catenin residues at its interfaces with the activating TCF/LEF transcription factors and the BCL9 co-activator are enriched in the Bottom 15% gate but depleted from the Top 10% gate. Conversely, sgRNA that truncate APC or introduce mutations at the β-TrCP-β-catenin interface are enriched in the Top 10% gate. This bird’s eye view of screen results confirmed that our strategy, based on measuring enrichment of each sgRNA in two gates designed to isolate cells with elevated or reduced WNT/β-catenin signaling strength, efficiently identified residues in BDC components required to mediate regulatory PPIs and post-translational modifications (PTMs).

To select missense mutations for deeper mechanistic analysis, we took a systematic approach to validating the results of our base editing screens (diagrammed in **Extended Data Fig.4a**). Preference was given to sgRNAs with z-scores ≥2 that targeted residues not mplicated in prior structural or biochemical studies and residues targeted by multiple different guides (**Extended Data Figs. 2 and 3**). Initial filtering yielded 370 guides across *CTTNB1* (80 sgRNAs), *AXIN1* (73 sgRNAs), *APC* (196 sgRNAs), and *GSK3B* (21 sgRNAs). Each sgRNA was individually re-tested for its effect on WNT/β-catenin signaling strength using the 7TS reporter in eHAP1 cells and, for those guides that showed a significant effect, a second validation test was conducted in HEK293T cells (**Extended Data Fig.4, Supplementary Table 2**). Most of the missense mutations that were successfully validated mapped to β-catenin and AXIN1 (**Extended Data Fig.4b-e**). The z-scores for missense mutations in APC and GSK3β were lower (2<z<3, **Extended Data Fig.3**) compared to β-catenin and AXIN1 (**Extended Data Fig.2**). During validation studies, these mutations in APC and GSK3β had only modest effects (**Extended Data Fig.4f and g**). We speculate that APC engages in redundant interactions that cannot easily be disrupted by point mutations and that inactivating mutations in GSK3β might impair cell growth. Regardless, we focused on the mechanistic analysis of mutations in β-catenin and AXIN1 for the remainder of the study.

### The sequence-function landscape of β-catenin

To identify new regulatory sites in the BDC, we first focused on analysis of missense mutations in β-catenin (**Fig.2a, 2b, Extended Data Fig.2a-2d**). For ease of description, we divided β-catenin into 6 regions (Sites 1-6) that mediate specific PPIs or regulatory functions (**Fig.2b, Extended Data Fig.5a**)^11^. Mapping Top 10% and Bottom 15% enrichment z-scores (above a cutoff of 2) onto a structural model^46,47^ revealed that our screen identified mutations in β-catenin located at known interaction interfaces with β-TrCP, BCL9, AXIN1, APC and the TCF/LEF transcription factors (**Fig.2a**). A detailed examination of structurally characterized PPI interfaces demonstrated that our screen identified key residues predicted to make contacts (defined as a distance <5 Å) between β-catenin and β-TrCP^44^ (**Extended Data Fig.5b**), β-catenin and BCL9^48^ (**Extended Data Fig.5c**), β-catenin and LEF1^49^ (**Extended Data Fig.5d**), and β-catenin and TCF4^50^ (**Extended Data Fig.5e**). Interestingly, we also identified mutations around these interfaces that were beyond the 5 Å distance cutoff for a contact, suggesting that allosteric effects or changes in local conformation can influence these interactions.

**Fig.2.**
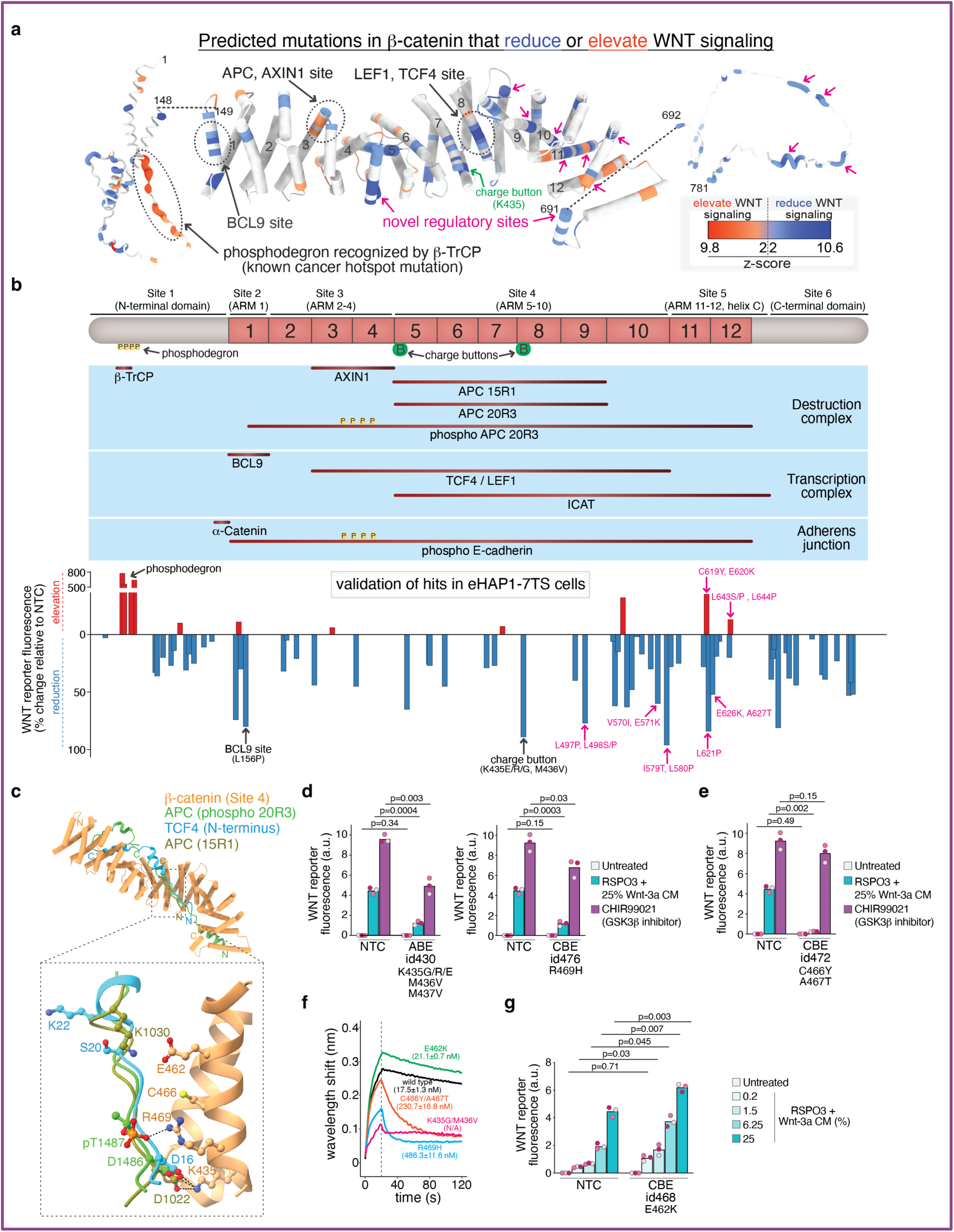
Base editing screens identify known and novel regulatory surfaces on β-catenin. **a,** A β-catenin backbone structural model (PDB 2Z6H) shows the position of missense mutations predicted (based on an editing window spanning N3-N10 of the protospacer) to be generated by sgRNAs with z-scores ≥ 2 in either the ABE or CBE screens^46^. Coloring denotes whether the predicted mutation increased (red) or decreased (blue) WNT/β-catenin signaling, based on whether the encoding sgRNA was enriched in the Top 10% or Bottom 15% gates used for sorting (see **1d**). Black arrows point to amino acid residues previously implicated in β-catenin function; magenta arrows highlight candidate residues that play a functional role in signaling. The 12 armadillo repeats of β-catenin are numbered on the last helix of each repeat. The N- and C-terminal segments of β-catenin (absent in published structures) were modeled by AlphaFold3 and are represented as worm models with a radius specified by the z-score^84^. **b,** Linear cartoon of β-catenin (top) showing Sites 1-6 and position of key charge button residues is aligned with a graphic showing regions of the ARM superhelix that interact with various partner proteins (brown lines in middle panel). The bar graph (bottom, aligned to the top cartoon) shows the position of sgRNAs that increased (red) or decreased (blue) WNT/β-catenin signaling strength (relative to a non-targeting control sgRNA, NTC) when individually introduced into eHAP1-7TS cells. Black arrows denote sgRNAs that target known β-catenin interaction interfaces and magenta arrows show sgRNAs selected for mechanistic analysis. **c,** Superimposed structural models showing the interaction of phosphorylated APC 20R3 (PDB 1TH1, green), TCF4 (PDB 1JDH, blue), and APC 15R1 (PDB 1JPP, olive) with β-catenin (PDB 2Z6H, sandy brown)^46,50,54,55^. In the zoomed view, a ball-and-stick depiction is used to show side chains of β-catenin residues identified as functionally significant in our screens. Electrostatic interactions are indicated with dashed black lines. **d,e,g,** WNT/β-catenin signaling strength (measured using the 7TS reporter, see **1c**) in HEK293T-7TS cells carrying various screen-identified mutations in β-catenin. Cells were treated with the indicated concentrations of Wnt-3a conditioned media (always in the presence of the potentiator RSPO3) or a GSK3β inhibitor (CHIR99021) for 24 h. Bars represent the mean of three biological replicates (*n*=3, corresponding points from each replicate are colored a different shade of pink). Each point represents the mean fluorescence from approximately 10,000 cells in a single biological replicate. Note that **d** and **e** show experiments that were conducted together (with the same control) but are depicted separately for clarity. Statistical significance was determined using multiple unpaired Student’s t-tests with Welch’s correction. P-values were adjusted using the two-stage step-up method (Benjamini, Krieger, and Yekutieli) to control the false discovery rate (FDR) at Q = 0.01. **f,** Biolayer interferometry (BLI) was used to assess association and dissociation kinetics between a GST-tagged TCF4 fragment (a.a. 1-53, immobilized on the BLI probe at 50 nM) and untagged, full-length β-catenin variants (1 µM). One representative curve is shown, with the affinity constant (*K_D_* ± s.d.) calculated from three independent replicates. See also Extended Data Figures 4 and 5 and Supplementary Table 2.

Overall, our screen and subsequent directed analysis in two different cell lines provided functional data on endogenous mutations at ∼186 residues in β-catenin (**Fig.2b, Extended Data Fig.5a**). While most β-catenin mutations reduced WNT signaling strength, we also identified gain-of-function mutations (e.g. phosphodegron mutations) that elevated signaling (**Fig.2b**). The high quality of these data is demonstrated by the identification of multiple residues that have been previously implicated in β-catenin function (highlighted in bold lettering in **Extended Data Fig.5a** and labeled in **Fig.2b**). Additionally, the tiling design of our libraries (**Extended Data Fig.1c**) allowed the identification of multiple sgRNAs that targeted the same residue, (**Extended Data Figs.2a-2d, 5a**). Our screens also identified unexpected mutations in β-catenin that either increase or decrease WNT/β-catenin signaling strength (but cannot be readily predicted by prior structural or biochemical studies). While we can explore only a small fraction of this dataset in this study, we hope that it serves as a valuable resource for the field to understand the function of β-catenin, a target for drugs being tested in CRC.

As an illustration of how scanning mutagenesis can be used to probe a structurally defined protein interaction interface, we focused on β-catenin Site 4, which forms an extended groove along the ARM superhelix between repeats 5-10 (**Fig.2b,2c**). The groove in Site 4 can bind (in a mutually exclusive manner based on structural data) to APC 20R and 15R repeats, TCF/LEF proteins, the negative regulator ICAT, and E-cadherin^49,51–57^ (**Fig.2c**). Our screen identified a column of mutations along one face of a helix that frames the Site 4 groove: K435G/M436G, R469H, C466Y/A467T, and E462K (**Fig.2c**). K435 is a well-characterized “charge button” residue known to be critical for multiple β-catenin interactions^11^. Indeed, sgRNAs targeting β-catenin K435 or a nearby basic residue R469 reduced signaling strength in response to WNTs or the GSK3-β inhibitor CHIR99021, consistent with their roles in anchoring β-catenin to TCF4^51,52^ (**Fig.2d**). Mutations at β-catenin C466/A467, implicated in nonpolar contacts with both APC and TCF4^58,59^, also markedly reduced signaling strength in response to WNTs (**Fig.2e**). β-catenin C466Y/A467T was fully active when signaling was initiated using CHIR99021, demonstrating these mutations did not compromise β-catenin integrity. Biolayer interferometry (BLI) binding assays using purified proteins confirmed that the R469H and C466Y/A467T mutations in β-catenin reduced its affinity for TCF4 by >10-fold (**Fig.2f**).

We also identified β-catenin mutations in this crowded PPI region that had effects only on specific interaction partners. E462K unexpectedly enhanced signaling strength in cells (**Fig. 2g**), suggesting selectively impaired interactions with the negative regulator APC. In binding assays with purified proteins, E462K did not change the affinity of the β-catenin-TCF4 interaction (**Fig.2f**). E462 likely forms an electrostatic interaction with K1030 in APC 15R1, though the K1030 side chain is disordered in the β-catenin-APC 15R-A crystal structure^54^ (**Fig.2c**). The residue at the position corresponding to APC K1030 in TCF4 points in the opposite direction (TCF4 K22) and thus cannot make a salt bridge with β-catenin E462, explaining why β-catenin E462K is a separation-of-function mutation that can discriminate between APC and TCF/LEF binding.

### New gain- and loss-of-function mutations in β-catenin

Several mutations in Sites 4-6 identified in our screen targeted residues not previously implicated in β-catenin function (sgRNAs highlighted in magenta lettering in **Fig.2b**, individual mutations shown in **Extended Data Fig.5a**). Mechanistic analysis was conducted in pooled HEK293T cell lines established after introduction of the base editor and specific sgRNA. Mutant allele frequencies in all cell lines were measured using Illumina sequencing to identify the dominant β-catenin mutation (**Extended Data Fig.6a**).

In β-catenin Site 4 (ARM5-10), three mutations (L497P/L498P, E571K and I579T/L580P, **Fig.3a**) reduced Wnt-3a-induced activation of both the 7TS reporter (**Fig.3b**) and transcription of the endogenous target gene *AXIN2* (**Extended Data Fig.7a**). Mutations in these residues impair signaling to the same extent as mutations that include the critical charge button (K435G/M436V), but they are not predicted to make contacts with any known interaction partners. To measure in-cell PPIs, β-catenin was immunoprecipitated (IP) from extracts of untreated cells or cells treated with Wnt-3a or CHIR99021 (**Fig.3c**). Importantly, these loss-of-function mutations did not decrease the abundance of β-catenin or abolish the interaction of β-catenin with AXIN1, APC or E-cadherin. However, these mutations decreased the interaction between β-catenin and multiple members of the TCF/LEF family of transcriptional activators. We propose that these mutations have allosteric or indirect effects selectively on the β-catenin-TCF/LEF interaction or that the published structures (which use N-terminal fragments of TCF/LEF) did not capture the full extent of this critical activating interaction.

**Fig. 3.**
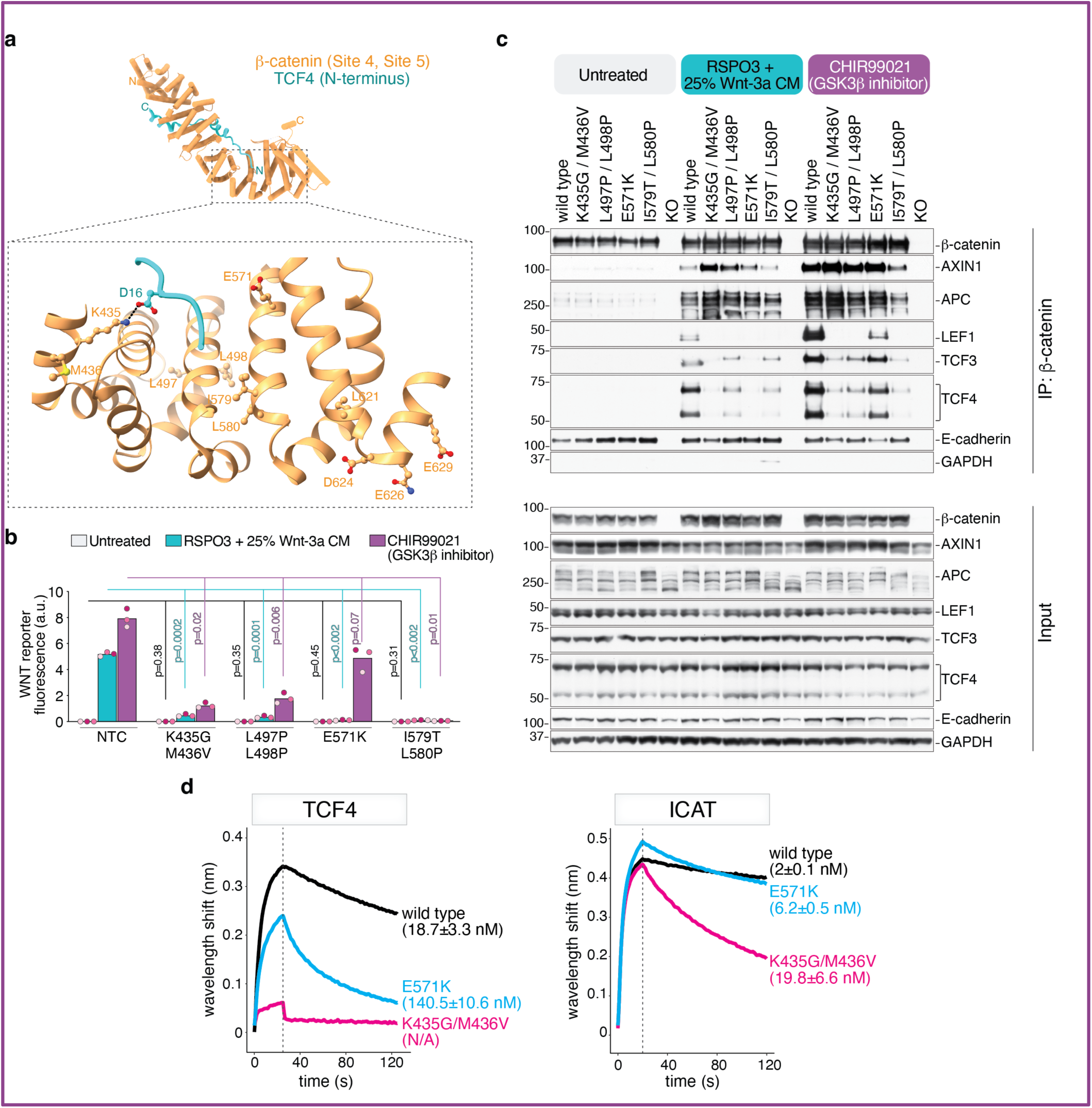
Mutations in β-catenin Site 4 affect interactions with the TCF/LEF transcription factors. **a,** Structure of β-catenin (PDB 2Z6H, sandy brown) in complex with the N-terminus of TCF4 (PDB 1JDH, blue)^50^. Zoomed view of Site 4 (ARM5-10) shows side chains of β-catenin residues identified as significant hits in our screen in ball-and-stick representation, including the key charge button K435 (see Fig. 2b). **b,** WNT/β-catenin signaling strength in HEK293T-7TS reporter cells carrying various mutations in β-catenin (confirmed by deep sequencing, **Extended Data Fig.6**) after treatment (24h) with RSPO3+Wnt-3a or CHIR99021. The bars represent the mean of three biological replicates (*n*=3, each in a different pink shade). Each data point is the mean 7TS fluorescence from ∼10,000 cells. Statistical significance was determined by Brown-Forsythe and Welch ANOVA followed by Dunnett’s T3 multiple comparisons test for pairwise comparisons between groups. **c,** Association of β-catenin with the indicated interaction partners assessed by co-immunoprecipitation (IP) from extracts of cell lines shown in **b** (*n*=2, one representative immunoblot shown). β-catenin knock-out (KO, *CTNNB1^-/-^*) cells serve as a specificity control for the IPs, association of β-catenin mutants with E-cadherin demonstrates protein integrity, and GAPDH is a loading control. **d,** BLI was used to assess association and dissociation kinetics between β-catenin variants and a GST-TCF4 fragment (left, same as **Fig.2f**) and GST-ICAT (right). All GST-tagged proteins were immobilized on the BLI probe at 50 nM and β-catenin variants were used at 1 µM (TCF4) and 0.5 µM (ICAT). One representative curve is shown, with the affinity constant (*K_D_* ± s.d.) calculated from three independent replicates. See also Extended Data Figures 5, 6 and 7.

The E571K mutation was distinguished by the property that it inhibited Wnt-3a-driven signaling but had little effect on signaling initiated by CHIR99021 (**Fig.3b**). Consistent with activity assays, the interaction between β-catenin E571K and TCF/LEF proteins was partially restored by CHIR99021 (**Fig.3c**). *In vitro* interaction assays (BLI) demonstrated that purified β-catenin E571K could still bind to purified TCF4, though its affinity was reduced by ∼7-fold relative to wild-type β-catenin (**Fig.3d**). The E571K mutation did not change the affinity between β-catenin and ICAT, a negative regulator that also binds in this region. In comparison, the β-catenin K435G/M436V charge button mutant had no detectable affinity for TCF4 and reduced affinity for ICAT (**Fig.3d**). Interestingly, E571K was the only mutant with a defect in Wnt-3a-induced accumulation of β-catenin (provisionally defined as the pool unphosphorylated at the N-terminal phosphodegron, **Extended Data Fig.7b**). Thus, the E571K mutation, in addition to impairing TCF/LEF binding, may cause a defect in BDC inactivation by WNTs.

We next turned to mutations in Site 5 (ARM11-12) that both decreased (**Fig.4a**, **Extended Data Fig.7c**) and increased (**Fig.4b**) WNT ligand sensitivity, without substantially altering protein abundance (**Fig.4c**). The reduction-of-function L621P and D624N/E626K/E628K mutations reduced the association between β-catenin and TCF/LEF proteins in cell lysates (**Fig.4c**), but did not affect Wnt-3a-induced accumulation of β-catenin (**Extended Data Fig.7d**). In purified binding assays, the D624K/E626K/E628K mutations reduced the affinity between β-catenin and both TCF4 and ICAT by ∼10-fold (**Fig.4f**). In comparison, both β-catenin mutations (C619Y/E620K and L643S/P, L644P) that increased the potency of Wnt-3a ligands also enhanced association with TCF4 in cell extracts (**Fig.4c**). This effect can be most clearly seen at low, subsaturating Wnt-3a concentrations (**Fig.4c**) and explains why cells carrying these mutations are hyper-responsive to WNTs (**Fig.4b**).

**Fig. 4.**
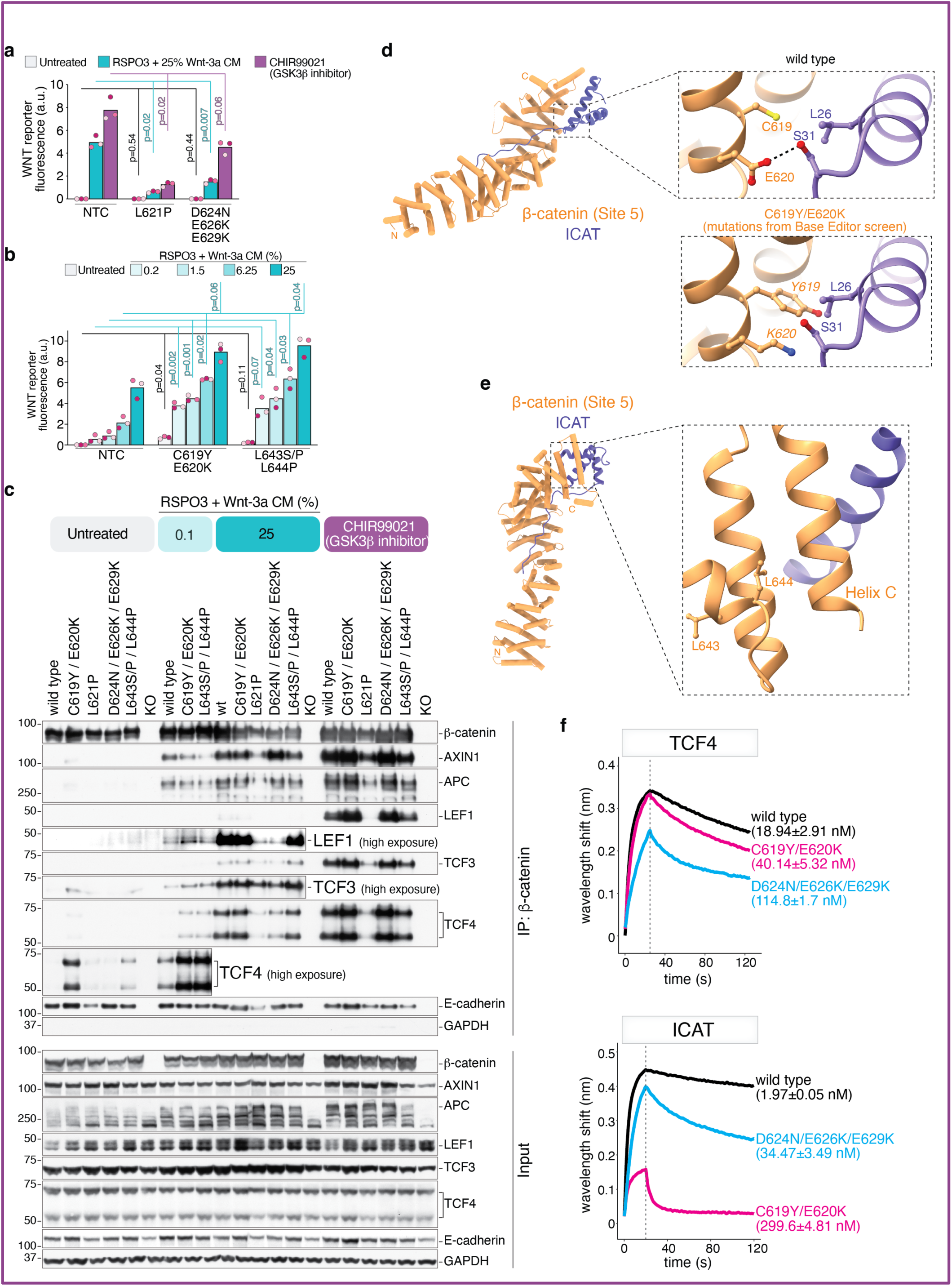
Mutations in β-catenin Site 5 (ARM 11-12) can reduce or enhance WNT/β-catenin signaling. **a,b,** WNT/β-catenin signaling strength in HEK293T-7TS reporter cells carrying mutations in β-catenin (confirmed by deep sequencing, **Extended Data Fig.6**) predicted to reduce (**a**) or enhance (**b**) WNT/β-catenin signaling. WNT/β-catenin signaling in cells carrying reduction-of-function mutations (**a**) was assessed after treatment with a saturating concentration of RSPO3+Wnt-3a or CHIR99021 (24h). Dose-response analysis (**b**) was used to assess sensitivity to Wnt-3a in HEK293T-7TS cells carrying putative gain-of-function mutations in β-catenin. The bars represent the mean of three biological replicates (*n*=3, each in a different pink shade). Each data point is the mean 7TS fluorescence from ∼10,000 cells. Statistical significance was determined by Brown-Forsythe and Welch ANOVA followed by Dunnett’s T3 multiple comparisons test for pairwise comparisons between groups. **c**, Association of β-catenin with the indicated interaction partners assessed by co-immunoprecipitation from extracts of cell lines shown in **a** and **b**, depicted as in **Fig.3c** (*n*=2, one representative immunoblot shown). **d,e,** Structural model of β-catenin (PDB 2Z6H, sandy brown) in complex with ICAT (purple) from PDB 1LUJ^46,56^. Zoomed region in **d** shows a potential steric clash created by the β-catenin C619Y/E620K gain-of-function mutation at the interface with ICAT. Zoom in **e** shows that the other gain-of-function mutations in β-catenin (L643S/P, L644P) are not directly at the PPI interface with ICAT. **f,** Association and dissociation kinetics between β-catenin variants and a GST-TCF4 fragment (top, same as **Fig.2f**) and GST-ICAT (bottom, same as **Fig.3d**). GST-tagged TCF4 and ICAT were immobilized on the BLI probe at 50 nM and β-catenin variants were used at 1 µM (TCF4) or 0.5 µM (for ICAT). Representative binding curves are shown, with the affinity constant (*K_D_* ± s.d.) derived from three biological replicates. See also Extended Data Figures 6 and 7.

The C619 and E620 residues are located in a region at the end of the β-catenin ARM superhelix that interacts with a helical bundle of the negative regulator ICAT (**Fig.4d**). Introduction of a bulky residue in C618Y or a charge reversal in E620K could disrupt this interface. In a purified binding assay, C619Y/E620K did not change the affinity of β-catenin with TCF4, but profoundly impaired the β-catenin-ICAT interaction (**Fig.4f**). Since ICAT and TCF4 bind to β-catenin in a mutually exclusive manner, the enhanced association of β-catenin C619Y/E620K with TCF/LEF factors in cell extracts (**Fig.4c**) is likely to be a secondary consequence of the impaired association of ICAT or other negative regulators^53,56,60^. The second activating mutation (β-catenin L643S/P, L644P) is not at the direct interface with ICAT but could potentially impact the interaction via an allosteric effect (**Fig.4e**). However, we were unable to produce this mutant protein in bacteria to measure affinities. This example further highlights the ability of base editing screens to identify useful separation-of-function mutations, especially important in analysis of proteins like β-catenin which have multiple interaction partners.

The isolation of mutations that can either increase or decrease WNT/β-catenin signaling strength in β-catenin ARM10-12 nominate this region as a largely unexplored regulatory site for the association of β-catenin with TCF/LEF transcription factors. Interestingly, a stapled peptide that binds in this region was shown to block the interaction between β-catenin and TCF4, but the functional effects on WNT/β-catenin signaling were not reported^61^. Several of these novel mutations in β-catenin also reduced the constitutively high level of WNT reporter activity in cancer cell lines carrying the CRC-driving APC^mcr^ oncoprotein (which cannot bind directly to AXIN1) (**Extended Data Fig.7e-h**). This activity profile is consistent with the model that these reduction-of-function mutations decrease WNT/β-catenin signaling downstream of the BDC at the level of the WNT transcriptional complex. Small molecules that bind in ARM10-12 may represent a useful strategy to disrupt the interaction between β-catenin and TCF4 in CRC.

### The sequence-function landscape of AXIN1

Base editing screens identified both gain- and loss-of-function missense mutations in AXIN1. Since AXIN1 is a negative regulator, gain-of-function mutations (**Fig.5a**, blue bars) suppress WNT/β-catenin signaling and loss-of-function mutations elevate WNT/β-catenin signaling (**Fig.5a**, red bars). Gain-of-function mutations in AXIN1 were considerably more common than those identified in β-catenin (**Fig.2b**, red bars) and were clustered in the DAX domain (**Fig.5a**, blue bars and **Extended Data Fig.2e-2h**).

**Fig. 5.**
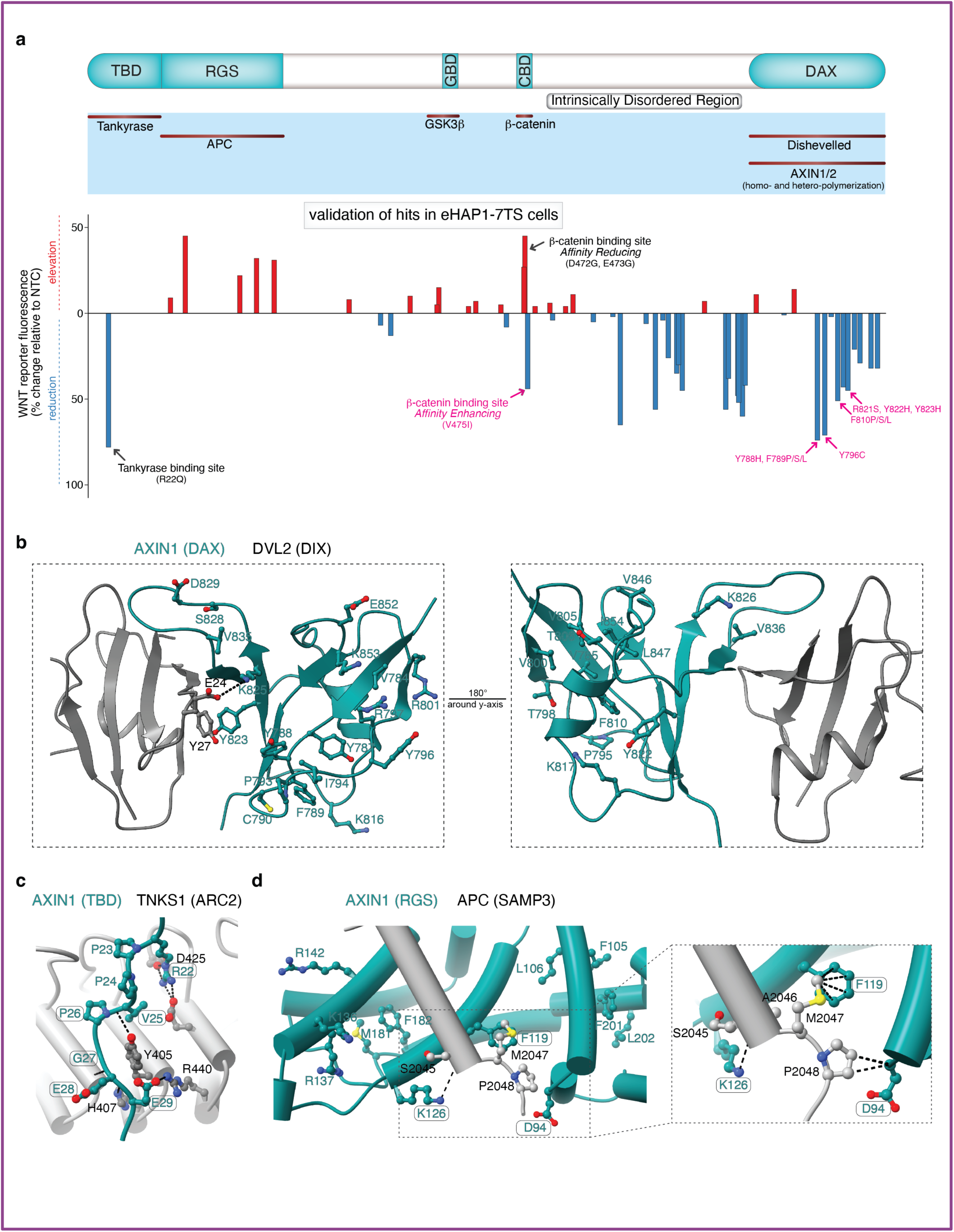
Base editing screens identify known and novel regulatory regions on AXIN1. **a,** Linear cartoon of AXIN1 (top) showing known regions that interact with partner proteins (middle). The bar graph (bottom, aligned to the cartoon at the top) shows the position of sgRNAs that increased (red) or decreased (blue) WNT/β-catenin signaling strength (relative to a non-targeting control sgRNA, NTC) when individually introduced into eHAP1-7TS cells. Black arrows denote sgRNAs that target known AXIN1 interaction interfaces and magenta arrows show sgRNAs selected for mechanistic analysis. TBD, Tankyrase Binding Domain; RGS, Regulator of G-Protein Signaling; GBD, GSK3β Binding Domain; CBD, β-catenin Binding Domain; DAX, Domain present in DVL and AXIN. **b,c,d,** Structural models showing PPI interfaces between various domains of AXIN1 (always in teal, defined in **a**) and the DIX domain of Dishevelled (DVL2, **b**)^67^, the Ankyrin repeat cluster 2 (ARC2) domain of Tankyrase 1 (TNKS1, **c**)^68^, and the SAMP3 repeat of APC (**d**)^19^. In **b**-**d**, ball-and-stick depiction is used to show side chains of AXIN1 residues predicted (located within the N3-N10 region of the protospacer) to be targeted by sgRNAs that were enriched (*z*-score of ≥ 2) in the Bottom 15% (**b**, **c**) or Top 10% (**d**) gates used for sorting (see **Fig.1d**). Rounded rectangles are drawn around residues previously identified as being critical for the depicted interaction and dotted lines show putative hydrophobic or electrostatic interactions. See also Extended Data Figures 2, 4 and 8.

The AXIN1 DAX domain has been implicated in two opposing roles^16,62–66^. DAX-mediated AXIN1 oligomerization is thought to increase the efficiency of β-catenin targeting by the BDC, thus keeping basal signaling low. However, in the presence of WNTs, the DAX domain mediates the recruitment of AXIN1 to DVL proteins at the receptor complex, ultimately resulting in inactivation of the BDC. Mutations enriched in the Bottom 15% gate were distributed throughout the DAX domain, including at the structurally-defined interface with the DIX domain of DVL^67^ (**Fig.5b**). Validation studies on a subset of these mutations revealed that they reduced Wnt-3a-induced reporter activation, but (in an important control) did not affect CHIR99021-induced signaling (**Extended Data Fig.8a)**. This reduction in WNT/β-catenin signaling strength was even observed for mutations (F810P/L, Y822H/Y823H) that reduced the abundance of AXIN1 (**Extended Data Fig.8b**). Surprisingly, none of these mutations in the DAX domain elevated basal signaling in the absence of Wnt-3a.

As with β-catenin, sgRNAs predicted to make mutations at known AXIN1 interaction interfaces were top hits in our screens (labeled in **Fig.5a**). Amongst the mutations enriched in the Bottom 15% gate, we identified several located at the known interface between AXIN1 and tankyrase enzymes (**Fig.5c, Extended Data Fig.2e-2h**). Tankyrases reduce AXIN1 abundance by poly-ADP-ribosylation-mediated ubiquitination^45,68^. These mutations (e.g. R22Q) increase AXIN1 abundance and consequently suppress WNT/β-catenin signaling (**Extended Data Fig.8**). Conversely, mutations at the interface between the AXIN1 RGS domain and Ser-Ala-Met-Pro (SAMP) motif on APC (**Fig.5d, Extended Data Fig.2g**) were highly enriched in the Top 10% gate, consistent with the prediction that they would increase WNT/β-catenin signaling strength by disrupting a key stabilizing PPI in the BDC^19^.

### Repairing an oncogenic BDC by affinity enhancing mutations in AXIN1

The fourth major region of AXIN1 highlighted by the mutations identified in our screens was the catenin binding domain (CBD), a peptide that binds to Site 3 in β-catenin as a short α-helix (**Extended Data Fig. 2h, Fig.6a**). This region was notable for mutations that both increased and decreased WNT/β-catenin signaling strength, a hallmark of regulatory sites in many proteins (**Fig.5a**). Two different mutations at the same residue (AXIN1 V475) had opposing effects on WNT/β-catenin signaling strength: V475A enhanced signaling while V475I diminished signaling (**Fig.6b, Extended Data 9a**). Binding assays using purified proteins revealed that the affinity of β-catenin for an AXIN1 CBD peptide carrying the V475I mutation was ∼2 fold higher (∼450 nM) compared to a WT CBD peptide (1 μM), while the affinity of AXIN1 V475A for β-catenin was markedly diminished (**Fig.6d**). This bi-directional correlation between affinity and activity suggested that quantitative changes in affinity between AXIN and β-catenin likely drive the observed cellular changes in WNT/β-catenin signaling strength.

**Fig. 6.**
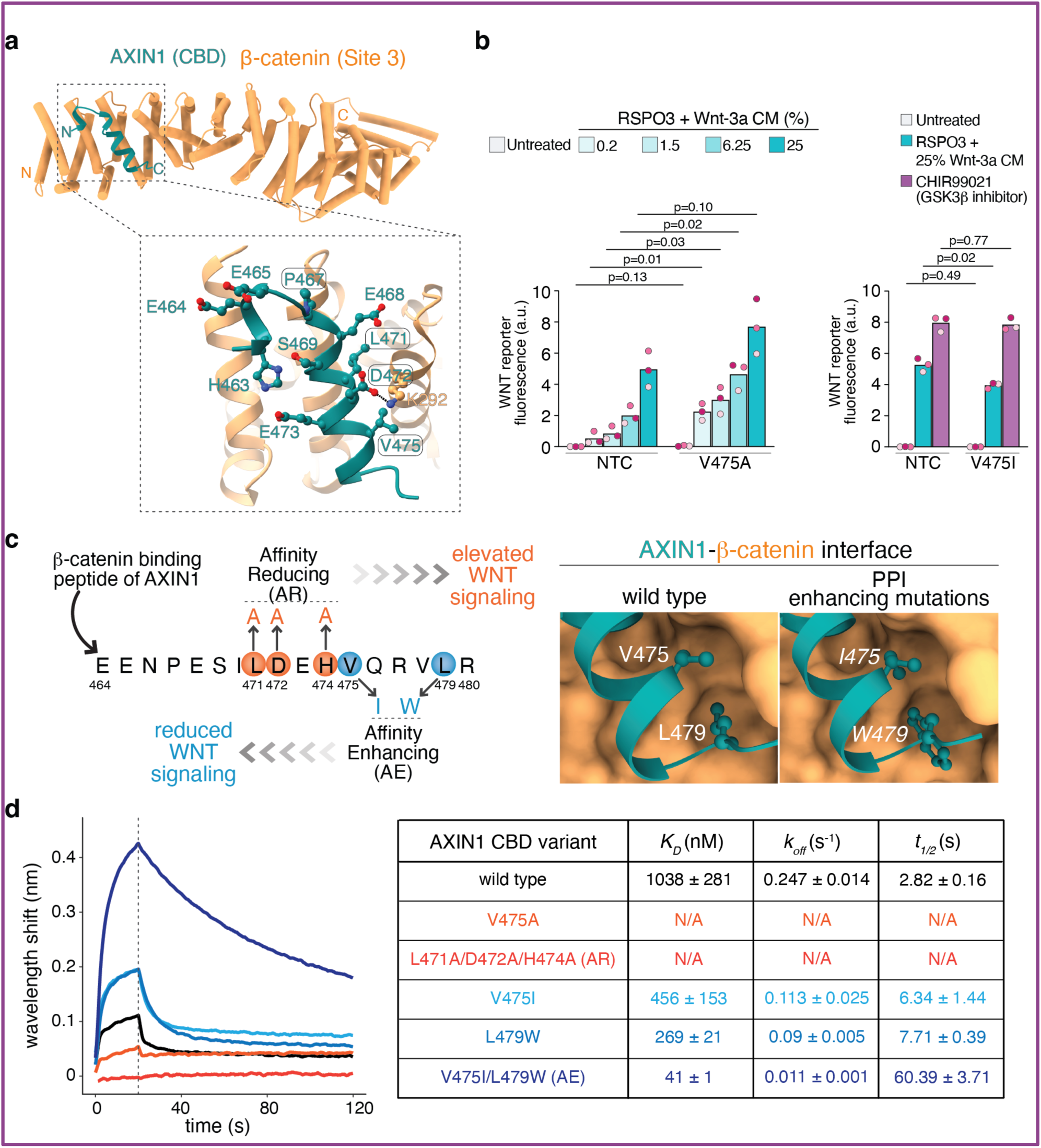
Base editing screens identify AXIN1 mutations that either increase or decrease its affinity for β-catenin. **a,** AlphaFold3 model of the AXIN1 CBD (green) bound in Site 3 (ARM3-4) of β-catenin (PDB 2Z6H, sandy brown). In the zoomed view of this largely hydrophobic interface, ball-and-stick depiction is used to show side chains of AXIN1 residues predicted (i.e. located within the N3-N10 region of the protospacer) to be targeted by sgRNAs that were enriched (*z*-score of ≥ 2) in the Top 10% gate used for sorting (see **Fig.1d**). Rounded rectangles are drawn around residues previously identified as being critical for the depicted interaction. **b,** WNT/β-catenin signaling strength in HEK293T-7TS reporter cells carrying mutations in AXIN1 (confirmed by deep sequencing, **Extended Data Fig.6**) predicted to enhance (left) or reduce (right) WNT/β-catenin signaling (assay identical to **Fig.4a,4b**). The bars represent the mean of three biological replicates (*n*=3, each in a different pink shade). Each data point is the mean 7TS fluorescence from ∼10,000 cells. Statistical significance was determined using multiple unpaired Student’s t-tests with Welch’s correction. P-values were adjusted using the two-stage step-up method (Benjamini, Krieger, and Yekutieli) to control the false discovery rate (FDR) at Q = 0.01. **c**, Mutations in the AXIN1 CBD peptide used to generate affinity-enhancing (AXIN1^AE^) and affinity-reducing (AXIN1^AR^) variants of AXIN1. **d**, *K_D_*, *k*_off_, and half-life (t_1/2_) values for purified β-catenin binding to variant GST-tagged AXIN1 CBD peptides shown in **b** and **c** were measured using BLI. GST-AXIN1 CBD peptides were immobilized at a concentration of 50 nM on the BLI probe and β-catenin was used at 1 µM in solution. Representative binding curves are shown, with the affinity constant (*K_D_* ± s.d.) derived from three biological replicates presented on the right.

This observation presented an opportunity to probe the AXIN1-β-catenin interaction in cells by tuning its affinity using mutations. Our goal was to compare the impact of these mutations on AXIN1-β-catenin affinity *in vitro* to their effect on WNT/β-catenin signaling and BDC integrity in cells. To further magnify affinity differences we combined V475I with L479W, identified in a phage display campaign to identify high-affinity β-catenin-binding stapled peptides^69,70^ (**Fig.6c**). The V475I/L479W double mutant AXIN1 CBD peptide demonstrated a 25-fold increase in affinity for β-catenin compared to the WT CBD (40 nM vs. 1 μM) in binding assays with purified proteins (**Fig.6d, Extended Data Fig.9b**). The increased affinity was almost completely driven by a ∼20-fold decrease in the rate constant for dissociation (*k*_off_) of β-catenin from an AXIN1 V475I/L479W CBD peptide (compared to a wild-type peptide). Thus, the half-life of a β-catenin-AXIN1 V475I/L479W complex is predicted to be ∼60 seconds, compared to ∼3 seconds for the wild-type complex. To disrupt the AXIN1-β-catenin interaction, we combined three mutations (L471A/D472A/H474A) in the AXIN1 CBD^71^ (**Fig.6c-d**) and established that this variant showed no detectable binding to purified β-catenin (**Fig.6d, Extended Data Fig.9b**). We hereafter refer to these mutations in AXIN1 as affinity enhancing (AXIN1^AE^: V475I/L479W) or affinity reducing (AXIN1^AR^: L471A/D472A/H474A) to distinguish them from wild-type AXIN1 (AXIN1^WT^) (**Fig.6c**).

To understand the consequences of increasing or decreasing the lifetime of the AXIN1-β-catenin complex in cells, we introduced the AE and AR mutations at the endogenous *AXIN1* locus in two isogenic HAP1 cell lines, one carrying an unmodified *APC* allele and a second with an oncogenic *APC^mcr^* allele (predicted to express an APC^mcr^ protein truncated at amino acid 1337 or APC 1337*) (**Extended Data Fig.10**). AXIN1 targeted co-IP experiments were used to assess BDC integrity. In cell lines expressing full-length APC, all three AXIN1 variants were associated with β-catenin and APC to a comparable extent under conditions of no or low Wnt-3a exposure (**Fig.7a, Extended Data Fig.9c**). Since AXIN1^AR^ has no measurable direct affinity for β-catenin (**Fig.6d**), it likely associates with β-catenin indirectly through APC^16,72^. In the presence of saturating Wnt-3a or CHIR99021 (both conditions that maximally inhibit GSK3β and activate signaling), AXIN1^AE^ pulled down much greater amounts of both β-catenin and APC compared to AXIN1^WT^ (**Fig.7a, Extended Data Fig.9c**). Thus, increasing the *in vitro* AXIN1-β-catenin affinity (**Fig.6d**) increased their association in cell only under conditions of high WNT exposure or GSK3β inhibition (both of which lead to high-amplitude signaling).

**Fig. 7.**
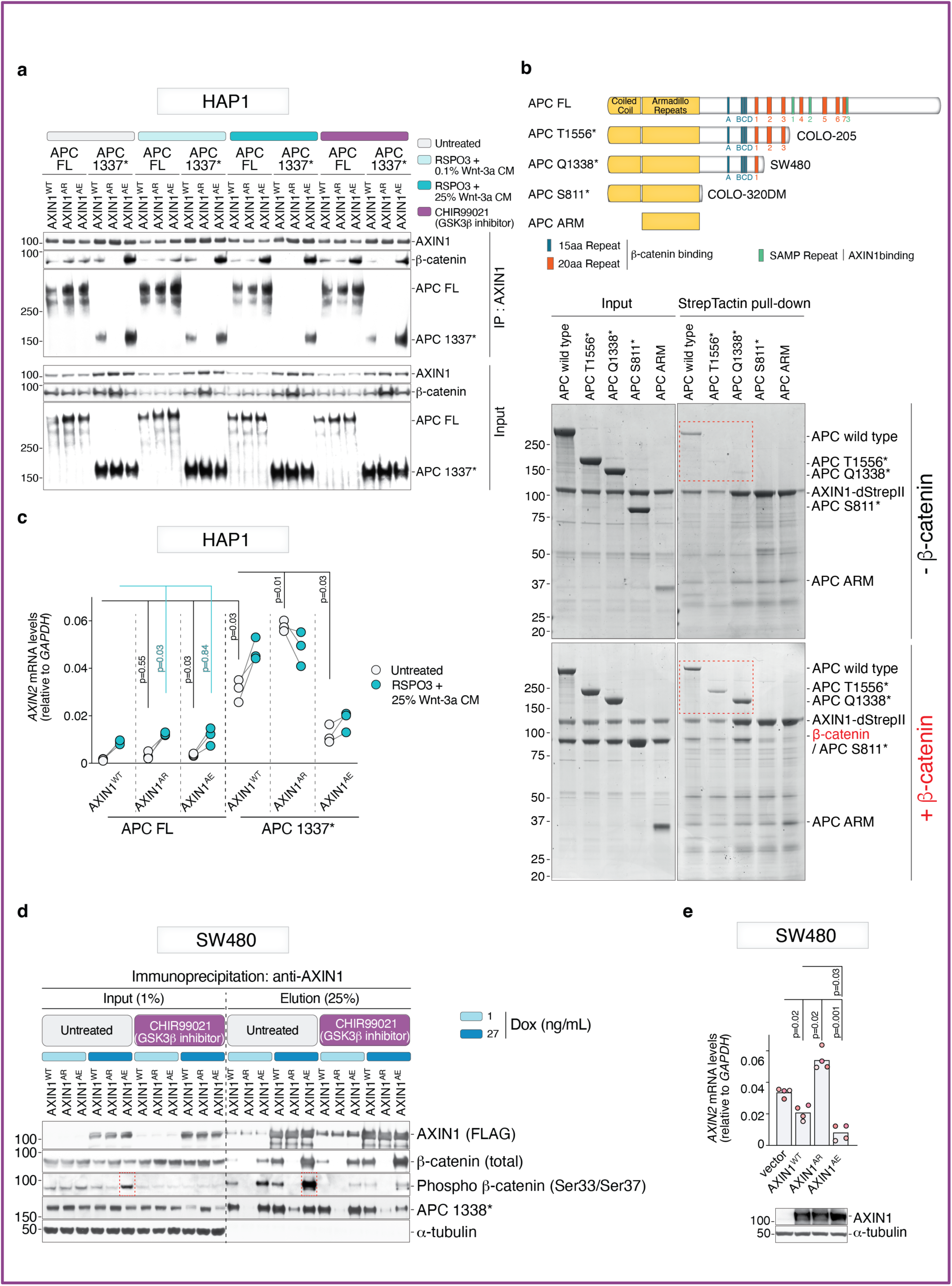
The AXIN1-β-catenin interaction is critical for the assembly of the oncogenic BDC. **a,** Association of endogenous AXIN1 with APC and β-catenin in extracts from clonal eHAP1 cells measured using an AXIN1 IP followed by immunoblotting. Sequence verified clonal cell lines (**Extended Data Fig.10**) contained the indicated combinations of full-length APC (APC FL) or truncated APC (APC 1337*) and one of three AXIN1 variants-- AXIN1^WT^, AXIN1^AE^ (V775I/L479W) and AXIN1^AR^ (L471A/D472A/H474A)-- with varying affinities for β-catenin (see **Fig.6d**). Extracts were made from untreated cells or cells treated (1h) with low-dose Wnt-3a, high-dose Wnt-3a or CHIR99021. **b,** A coomassie-stained gel depicts the association between purified, Strep II-tagged AXIN1 (bait) and purified APC variants (schematically shown above the gel) in the absence (top gel) or presence (bottom gel) of purified β-catenin. APC^mcr^ variants only associate with AXIN1 in the presence of β-catenin (dotted red boxes). **c,** The abundance of *AXIN2* mRNA, encoded by a WNT target gene, was used to measure the strength of signaling in response to Wnt-3a in HAP1 cells lines carrying the indicated combination of *APC* and *AXIN1* alleles (described in **a**). Three independent clonal cell lines were used for each *AXIN1* and *APC* allelic combination, with a line used to link data points from the same cell line. **d,** Association of doxycycline (Dox)-inducible, FLAG-tagged AXIN1 variants (AXIN1^WT^, AXIN1^AE^ or AXIN1^AR^) with endogenous truncated APC (APC 1338*) and endogenous β-catenin was measured using a FLAG IP from extracts of untreated or CHIR99021 treated SW480 CRC cells. Dotted red boxes show that expression of AXIN1^AE^ increases the abundance of phosphorylated β-catenin in both cell extracts and BDC IPs. **e,** WNT/β-catenin signaling strength in SW480 CRC cells stably expressing FLAG-tagged AXIN1 variants. The bars represent the mean of four biological replicates (*n*=4, each in a different pink shade). Each data point represents the mean *AXIN2* mRNA abundance (relative to a control housekeeping mRNA) from three technical replicates. In panels **c** and **e**, statistical significance was determined by Brown-Forsythe and Welch ANOVA followed by Dunnett’s T3 multiple comparisons test for pairwise comparisons between groups. Experiments in **a**, **b**, and **d** were repeated at least twice (*n*=2) and one representative gel or immunoblot is shown. See also Extended Data Figures 9 and 10.

In *APC^mcr^* cells, which express a truncated APC (APC 1337*) protein that lacks all three of the AXIN1-binding SAMP repeats, AXIN1^AE^ bound much greater amounts of β-catenin and APC compared to AXIN1^WT^ under all conditions tested. Conversely, the interaction between AXIN1^AR^ and both β-catenin and APC was abolished. Thus, in the absence of the RGS-SAMP mediated interaction between AXIN1 and APC^mcr^, β-catenin can bridge the two proteins by simultaneously binding to APC^mcr^ (through its 15R A-D repeats) and AXIN1 (through its CBD)^16,72^. Reconstitution experiments using purified proteins support the idea that β-catenin functions as the linchpin of the oncogenic BDC^16^ (**Fig.7b**). Oncogenic APC variants truncated before the SAMP repeats only associated with AXIN1 in the presence of β-catenin. Further truncations of APC that remove the 15R repeats abolished the capacity of β-catenin to bridge APC and AXIN1 (**Fig.7b**).

There was a notable difference in WNT/β-catenin signaling behavior, as measured by *AXIN2* mRNA abundance, between cell lines expressing full-length APC or truncated APC (APC 1337*). In the context of full-length APC, basal and Wnt-3a-induced signaling in cell lines carrying any of the three AXIN1 variants was comparable. However, the results were different in APC^mcr^ cells: AXIN1^AR^ led to a further increase in *AXIN2* mRNA levels and nuclear β-catenin abundance while the AXIN1^AE^ variant had the opposite effect (**Fig.7c**, **Extended Data Fig.9d**). Thus, cells carrying an oncogenic *APC^mcr^* allele were more sensitive to changes in the affinity between AXIN1 and β-catenin than those carrying WT APC.

To test this concept in a cancer cell line, we stably expressed AXIN1^WT^, AXIN1^AE^ or AXIN1^AR^ in the human CRC cell line SW480 using an inducible expression system. SW480 carries a mutant *APC* allele that encodes a protein truncated at amino acid residue 1338 (nearly identical to the APC 1337* protein expressed in our engineered HAP1 cells). SW480 cells have elevated WNT/β-catenin signaling and *AXIN2* expression due to impaired BDC function, leading to a defect in β-catenin phosphorylation and degradation^73^. In co-IP experiments, AXIN1^AE^ showed enhanced association with β-catenin, while AXIN1^AR^ failed to associate with β-catenin (consistent with HAP1 results) (**Fig.7d**). More importantly, β-catenin phosphorylated at its N-terminal degron motif (and thus marked for degradation) was considerably more abundant in cells expressing AXIN1^AR^, showing that enhancing the affinity of AXIN1 and β-catenin can improve the activity of an oncogenic BDC complex (**Fig.7d**). Accordingly, *AXIN2* mRNA abundance, a metric of WNT/β-catenin signaling strength, was suppressed to a greater extent in SW480 cells expressing AXIN1^AE^ compared to cells expressing AXIN1^WT^. Interestingly, WNT/β-catenin signaling was hyperactivated in SW480 cells expressing AXIN1^AR^, likely because replacement of the endogenous WT AXIN1 with AXIN1^AR^ resulted in disassembly and complete inactivation of the BDC (**Fig.7e**). Taken together, these data show that the AXIN1-β-catenin interaction is a rate-limiting step in oncogenic WNT/β-catenin signaling that can be manipulated to increase or decrease the activity of the impaired BDC found in cells that carry *APC^mcr^* alleles (**Fig.7e**).

## Discussion

In this work, base editing screens provided mechanistic insights into the BDC, a complex signaling machine that functions as the nerve center of WNT/β-catenin signaling, one of the most important developmental and stem cell signaling systems that also drives the majority of human CRCs. A longstanding, unresolved challenge in WNT signalling is understanding the biochemical differences between the wild-type, ligand-regulated BDC and the oncogenic BDC, hobbled by truncating mutations in APC. Human cancer genetics first revealed that oncogenic WNT/β-catenin signaling is not driven by the complete loss of APC function, as might be expected if it were a simple gatekeeper^37,74^. Instead, at least one *APC* allele in both sporadic and familial CRC cases is truncated such that it loses all three SAMP repeats required for its interaction with AXIN1 but retains at least one of the 20R repeats that can mediate a high affinity interaction with β-catenin (**Fig.1b, Extended Data Fig.7e**). This arrangement results in an intermediate level of ligand-independent WNT/β-catenin signaling that drives tumorigenesis but avoids senescence or cell death^74,75^. Understanding the detailed biochemistry of how the oncogenic BDC calibrates this just-right level of signaling flux also is central to therapeutic development. Given the pervasive role of WNT/β-catenin signaling in tissue homeostasis, a clinically viable WNT inhibitor must differentiate between mutation-driven oncogenic signaling and ligand-driven physiological signaling.

We propose that BDC assembly and regulation is fundamentally different in cells expressing full-length APC and APC^mcr^. These insights emerged from experiments that allowed us to increase or decrease the affinity of a specific PPI, the AXIN1-β-catenin interaction, using mutations at endogenous gene loci (thus avoiding the potentially confounding consequences of overexpression). In the presence of full-length APC, increasing or decreasing the AXIN1-β-catenin affinity across a >50-fold range above and below the native *K_D_* (∼1 μM), had very little effect on BDC assembly or target gene induction under basal, ligand-free conditions. This is likely because the APC 20R repeats are phosphorylated by GSK3β under basal conditions, drastically increasing their affinity (*K_D_* ∼1 nM *in vitro*) for the same groove (ARM 3-4) in β-catenin occupied by the AXIN1 CBD (**Fig.8a**)^55,58^. Thus, β-catenin is not available to interact with even the high affinity (*K_D_* ∼40 nM) AXIN1^AE^ variant used in our experiments. Essentially, the phosphorylation of APC by GSK3β makes the AXIN1-β-catenin interaction dispensable for β-catenin recruitment into the BDC or for the assembly of the BDC itself (**Fig.8b**). Consistent with this view, when GSK3β is inhibited either by CHIR99021 or by high Wnt-3a addition, the AXIN1-β-catenin affinity becomes relevant: AXIN1^AE^ pulls down much more β-catenin and APC (**Fig.7a**). Under these conditions, APC phosphorylation drops, reducing the affinity of the 20R repeats for β-catenin by ∼300-1000-fold and thus vacating the ARM3-4 groove for the AXIN1 CBD (**Fig.8c**)^17,18^.

**Fig. 8.**
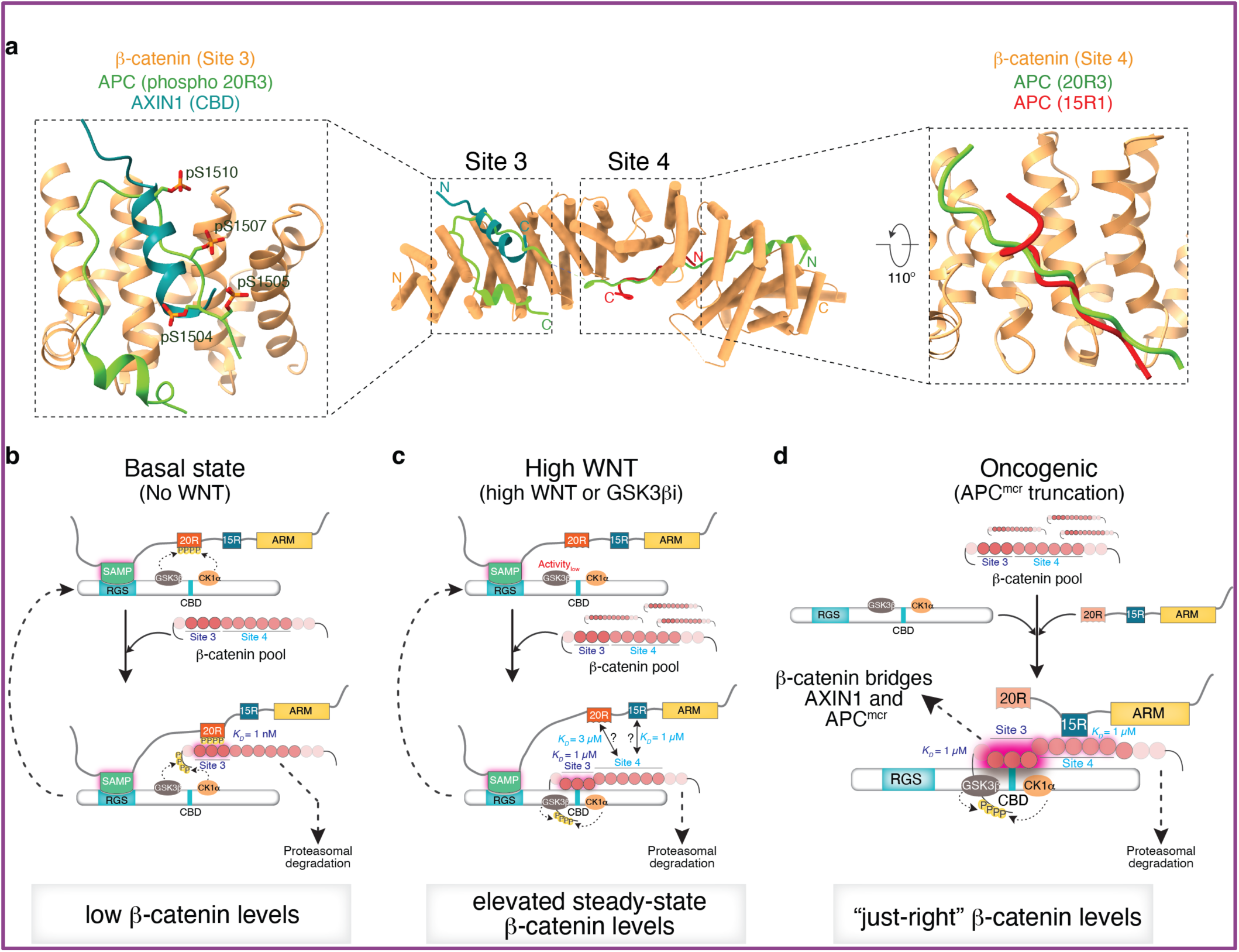
Different modes of β-catenin recognition by the wild-type and oncogenic BDC. See text for details. **a,** Structural superposition shows that the AXIN1 CBD (PDB 1QZ7) and phosphorylated APC 20R (p20R; PDB 1TH1) both bind to overlapping (and hence mutually exclusive) surfaces in β-catenin (PDB 2Z6H) Site 3, though the affinities are vastly different (*K_D_*∼1 nM for APC p20R and *K_D_*∼1 μM for AXIN1)^17,18,46,58,71^. APC 15R motifs (PDB 1JPP) and the extended regions of 20R motifs (PDB 1TH1) can also bind in β-catenin (PDB 2Z6H) Site 4, but AXIN1 only binds in Site 3^46,54,58^. **b,** In the presence of full-length APC, APC and AXIN1 can associate directly, allowing phosphorylation of the APC 20R motifs by the AXIN1-bound kinases and thus creating a high affinity (*K_D_*∼1 nM) binding site for β-catenin Site 3. In this regime, AXIN1 cannot directly bind to β-catenin, and the steady state concentration of β-catenin is kept low (limited by the APC p20R-β-catenin affinity). **c,** Inhibition of GSK3β by WNTs or CHIR99021 reduces APC phosphorylation, thus sharply decreasing the affinity of 20R motifs for β-catenin Site 3. Thus, β-catenin is recruited into the BDC primarily through its moderate affinity interaction (*K_D_*∼1 μM) with AXIN1 (or APC via Site 4), which requires a higher steady state β-catenin concentration for significant association. **d,** When APC is truncated in CRC, β-catenin becomes the linchpin, recruiting AXIN1 and APC via moderate affinity interactions with Site 3 and Site 4, respectively. Unlike the case where APC is intact (**b**, **c**), there are no cycles of β-catenin recruitment and degradation; each oncogenic BDC disassembles when β-catenin is phosphorylated and degraded. The steady-state β-catenin level is autoregulated-- an increase in the β-catenin concentration increases BDC assembly and hence enhances the degradation rate while a decrease reduces BDC assembly and hence decreases the degradation rate.

In cells with APC^mcr^, the AXIN1-β-catenin interaction becomes a critical determinant of BDC assembly for two key reasons: APC^mcr^ cannot bind directly to AXIN1 because it has lost its SAMP motifs and, as a consequence, the remaining 20R repeats in APC^mcr^ cannot be phosphorylated by GSK3β to generate a high-affinity β-catenin binding site (**Fig.8d**). Indeed, point mutations at the interface between AXIN1 CBD and β-catenin ARM 3-4 (AXIN1^AR^) prevented the association between AXIN1 and both β-catenin and APC^mcr^. Our data support the model that β-catenin serves as the linchpin of the oncogenic BDC, bridging AXIN1, which binds to ARM 3-4 via its CBD, and APC^mcr^, which binds to the extended ARM5-9 region via its 15R repeats (**Fig.8d**). Accordingly, the interaction of purified AXIN1 with APC^mcr^ depends on the presence of β-catenin. Full-length APC binds to AXIN1 regardless of the presence of β-catenin because its SAMP motifs can directly engage the RGS domain of AXIN1. Conversely, more extensive APC truncations that lack the 15R repeats fail to associate with AXIN1 regardless of β-catenin (**Fig.7b**). These data are consistent with prior work showing that disruption of β-catenin in cells prevents the co-precipitation of AXIN1 and truncated APC from cell extracts^72^.

We propose that β-catenin becomes both a substrate and a key assembly factor for the oncogenic BDC, thus promoting its own degradation and limiting its own abundance to achieve the just-right signaling flux necessary for CRC development. This mode of autoregulation is conceptually analogous to substrate-assisted catalysis, where the concentration and affinity of the substrate (β-catenin) for the enzyme (BDC) determines catalytic efficiency. Thus, steady-state β-catenin abundance in CRC cells is self-regulated by its own availability and critically dependent on its affinities for AXIN1 and APC^mcr^. In support of this idea, increasing or decreasing the AXIN1-β-catenin affinity has a corresponding effect on the abundance of phosphorylated β-catenin, the immediate product of the reaction catalyzed by the BDC (**Fig.7d**). Changes in the affinity between β-catenin and APC^mcr^ would also be predicted to have analogous effects on the catalytic efficiency of the oncogenic BDC, though such mutations were not identified in our screens. In the presence of full-length APC, β-catenin is simply recruited as a substrate to the BDC, which is optimized for suppressing basal signaling by minimizing β-catenin abundance (**Fig.8a**).

The substrate-assisted model of the oncogenic BDC has therapeutic implications. Small molecule drugs that slow the first-order dissociation of β-catenin from AXIN1 or APC^mcr^ (thus mimicking of the affinity-enhancing mutations identified in our screens) may partially repair the activity of the oncogenic BDC (without significantly affecting the native BDC). Similarly, drugs that disrupt these interactions should super-activate WNT/β-catenin signaling, also a detrimental outcome given the just-right model for CRC development^74,75^. Substrate affinity and substrate turnover can be competing properties that would need to be balanced in any such strategy. If the affinity between β-catenin and the oncogenic BDC is too high, product inhibition may limit BDC activity if phosphorylated β-catenin cannot be efficiently ubiquitinated or extracted for proteasomal degradation.

We finish by noting that base editing screens, which now enable the installation and functional evaluation of point mutations across endogenous proteins, have distinct advantages over loss-of-function and overexpression screens. Beyond gene identification, genetic screens in model organisms have provided insights into biochemical mechanisms of protein function by enabling the isolation of alleles that carry mutations at one or a small number of amino acid residues. These point mutations, often on protein surfaces, can identify enzyme active sites, protein-protein or protein-ligand interaction interfaces, and allosteric sites. The most informative (but rare) alleles are gain-of-function and separation-of-function alleles. Gain-of-function alleles can identify positive or instructive (rather than permissive) functions of a protein, identify rate limiting steps, and bypass genetic redundancy. We identified several such mutations in both β-catenin and AXIN1: mutations that increased the in-cell association of β-catenin with TCF/LEF proteins and mutations in the CBD and C-terminal disordered region of AXIN1 that suppress WNT/β-catenin signaling. A key signature of a regulatory site, identified both for the AXIN1 CBD and the ARM 11-12 region of β-catenin, are distinct mutations that have opposite effects on signaling strength. Second, separation-of-function mutations are especially valuable in mapping sequence-function relationships in multi-functional proteins (like β-catenin). Even in the very congested PPI interfaces on β-catenin, our screens identified mutations that distinguished between interactions with APC, TCF/LEF, and ICAT. Finally, our analysis of residues at known interfaces between β-catenin and its binding partners showed that the functional impact of mutations at endogenous genes can sometimes be difficult to predict from purely structural proximity considerations or *in vitro* binding assays. Hence, *in situ* base-editing screens can be used for the in-cell functional annotation of PPI interfaces identified by structural and biochemical studies.

## Methods

### Reagents and antibodies

Wnt-3a conditioned medium was produced as described previously^29^. Recombinant human R-Spondin 3 (RSPO3) was purchased from R&D Systems (Cat#3500-RS-025). CHIR99021 was purchased from Cayman Chemical (Cat#13122). The following primary antibodies were used for immunoblotting and immunofluorescence: β-catenin (BD Biosciences, Cat#610154, 1:3,000 for immunoblotting), Non-phospho (Active) β-Catenin (Ser33/37/Thr41) (Cell Signaling Technology, Cat#8814, 1:1,000 for immunoblotting, 1:200 for immunofluorescence), Phospho β-Catenin (Ser33/37) (Cell Signaling Technology, Cat#2009, 1:1000 for immunoblotting), AXIN1 (Cell Signaling Technology, Cat#2074, 1:1,000 for immunoblotting), APC (Millipore-Sigma, Cat#OP44, 1:200 for immunoblotting), LEF1 (Cell Signaling Technology, Cat#2230, 1:1,000 for immunoblotting), TCF3 (Cell Signaling Technology, Cat#2883, 1:1,000 for immunoblotting), TCF4 (Cell Signaling Technology, Cat#2569, 1:1,000 for immunoblotting), E-Cadherin (Cell Signaling Technology, Cat#3195, 1:1,000 for immunoblotting), GAPDH (Proteintech, Cat#60004-1-Ig, 1:10,000 for immunoblotting), α-Tubulin (Proteintech, Cat#66031-1-Ig, 1:10,000 for immunoblotting), and GST (Bethyl Laboratories, Cat#A190-122A, 1:1000 for immunoblotting). The following secondary antibodies were used: Peroxidase conjugated donkey anti-mouse/rabbit IgG (Jackson ImmunoResearch), IRDye 800CW donkey anti-mouse/rabbit IgG (LICORbio), and Alexa Fluor 488 conjugated donkey anti-mouse/rabbit IgG (Thermo Fisher Scientific).

### Cell culture

HAP1, a near haploid human cell line and eHAP1, a fully haploid human cell line^32^, were gifts from Jan Carette, Stanford University. These cell lines were authenticated by morphology and ploidy analysis as described below to confirm identity. HEK293T, RKO, DLD-1, and SW480 were obtained from American Type Culture Collection (ATCC) and their identity was authenticated by the vendor using STR profiling. All cell lines were confirmed to be mycoplasma-free using an isothermal PCR-based assay (InvivoGen, Cat#rep-mys-10). Experiments were performed using cells between passages 3 and 25. HAP1 and eHAP1 cells were grown in Iscove’s Modified Dulbecco’s Medium (IMDM) containing L-Glutamine (cytiva, Cat#SH30243FS) and supplemented with 10% fetal bovine serum (FBS) (MilliporeSigma, Cat #F0926), extra glutamine (2 mM) (cytiva, Cat#SH3003401), and penicillin-streptomycin (50 U/mL) (Thermo Fisher Scientific, Cat #15140163). HEK293T, RKO, DLD-1, and SW480 cells were grown in Dulbecco’s Modified Eagle Medium (DMEM) containing high glucose, with 4.0mM L-glutamine, and sodium pyruvate (cytiva, Cat #SH30243) and supplemented with 10% FBS, 1x MEM non-essential amino acids solution (Thermo Fisher Scientific, Cat #11140076), and penicillin-streptomycin (50 U/mL). Cells were subcultured every 3 days at a splitting ratio of 1:10 using 0.25% Trypsin-EDTA (Thermo Fisher Scientific, Cat #25200072) for HAP1 and eHAP1 cells or 0.05% Trypsin-EDTA (Thermo Fisher Scientific, Cat# 25300062) for HEK293T, RKO and DLD-1 cells. All cell lines were cultured in a humidified atmosphere containing 5% CO2 at 37 °C. HAP1 and eHAP1 cells were routinely tested for ploidy state every fifth passage by flow cytometry with propidium iodide staining as described previously^76^.

### Generation of WNT transcriptional reporter cell lines

Clonal WNT transcriptional reporter cell lines were generated in eHAP1, HEK293T, RKO, and DLD-1 cells. These lines contain a fluorescent reporter construct downstream of a WNT-responsive element. For RKO and DLD-1 lines, the lentiviral plasmid 7xTcf-eGFP (7TG) was used. This plasmid contains seven repeats of an optimized TCF/LEF binding site driving expression of eGFP. It was generated from 7xTcf-eGFP//SV40-PuroR^77^ by digestion with BsrGI and SalI to remove the SV40-PuroR cassette, creating a selection marker-free backbone that is compatible with CRISPR base editor plasmids. For eHAP1 and HEK293T lines, the 7xTcf-mScarlet (7TS) lentiviral plasmid was used, in which the eGFP cassette was replaced with the coding sequence for mScarlet. Lentivirus was produced by co-transfecting the respective reporter plasmid (7TG or 7TS) with the packaging plasmids psPAX2 (Addgene Plasmid #12260) and pMD2.G (Addgene Plasmid #12259) into 293FT (Thermo Fisher Scientific) cells using polyethylenimine (PEI; Polysciences, Inc., Cat# 23966-2). Supernatant containing lentivirus was collected at 48 h post-transfection and filtered (0.45 µm). Recipient clonal cell lines were transduced at an MOI of ∼0.5 in the presence of polybrene (8 µg/mL; Millipore-Sigma, Cat#H9268). Two days post-transduction, cells were treated with 50% Wnt-3a conditioned medium for 24 h followed by FACS to isolate eGFP/mScarlet positive single cells using a SH800S cell sorter (Sony Biotechnology) into 96-well plates. Multiple individual clones (∼24) were picked and expanded for each reporter line. Reporter activity was validated in expanded clonal lines after 24 h treatment with 50% Wnt-3a conditioned medium. Reporter fluorescence (eGFP or mScarlet) was measured by flow cytometry using a 488 nm laser and FL2 525/50 nm bandpass filter (eGFP) or a 561 nm laser and FL3 600/60 nm bandpass filter (mScarlet). Clonal lines with optimal growth rate, morphology, and a high dynamic range of reporter fluorescence were selected for downstream assays. High dynamic range was defined as achieving at least 100-fold induction of reporter fluorescence upon Wnt-3a stimulation compared to control.

### FACS-based pooled CRISPR Base Editing Screens

#### Library construction

For tiled sgRNA library design, we used the Beagle tool from the Broad Institute Genetic Perturbation Platform. The resulting library was filtered to exclude guide sequences containing *BsmBI* restriction sites. The Ensembl transcript IDs used for guide design were: *CTNNB1* (ENST00000349496), *APC* (ENST00000257430), *AXIN1* (ENST00000262320), and *GSK3B* (ENST00000264235). *BsmBI* recognition sites and appropriate overhangs were appended to each targeting sequence sequence for Golden Gate cloning into base editor plasmids (pRDA 429, Addgene#179098 expressing ABE8e Cas9-NG and pRDA 336, Addgene #179095 expressing BE3.9 Cas9-NG). Primer sites were included for amplification of subsets from the same synthesis pool. The final oligonucleotide sequence was as follows 5’-[Forward Primer]*CGTCTCACACCG*[targeting sequence, 20 nt]*GTTTCGAGACG*[Reverse Primer]-3’. Oligonucleotide pools were synthesized by CustomArray. PCR amplification of the oligonucleotide pools used 25 µL of 2x NEBNext PCR Master Mix (New England Biolabs, Cat# M0541S), 2 µL of the oligonucleotide pool (∼2 ng), 5 µL of primer mix at a final concentration of 0.5 µM, and 18 µL of nuclease-free water. PCR cycling conditions were: (1) Initial denaturation: 98 °C for 1 min; (2) 98 °C for 10 s; (3) 62 °C for 10 s; (4) 72 °C for 20 s; (5) cycle through steps 2-4 for 18 cycles; (6) Final extension: 72 °C for 2 min. The following primer (5’-3’) pairs were used for amplification:

APC-ABE: Forward: GGTCGTCGCATCACAATGCG, Reverse: TCTCGAGCGCCAATGTGACG

APC-CBE: Forward: GTGTAACCCGTAGGGCACCT, Reverse: GTCGAGAGCAGTCCTTCGAC

CTNNB1-ABE: Forward: CATGTTGCCCTGAGGCACAG, Reverse: CCGTTAGGTCCCGAAAGGCT

CTNNB1-CBE: Forward: AGGCACTTGCTCGTACGACG, Reverse: ATGTGGGCCCGGCACCTTAA

AXIN1-ABE: Forward: GTCGCTAGCTAGGTCCGAAC, Reverse: TGTGCTGCGCAAGACCACGT

AXIN1-CBE: Forward: CAGCGCCAATGGGCTTTCGA, Reverse: AGCCGCTTAAGAGCCTGTCG

GSK3B-ABE: Forward: ACATGGCGCTGTCCCAGATG, Reverse: AACCTACGCCATGGTGCGTG

GSK3B-CBE: Forward: CTACAGGTACCGGTCCTGAG, Reverse: GTACCTAGCGTGACGATCCG

Control: Forward: TTACTAGACTGCGGGGCCAC, Reverse: CGGCTCGATCTGCGACGAAT

The resulting amplicons were resolved on a 2% TAE gel and purified by Zymoclean Gel DNA Recovery Kit (Zymo Research, Cat#D4008) and ligated into the pre-digested base editor plasmids using Esp3I (Thermo Fisher Scientific, Cat# ER1281) and T4 DNA ligase (New England Biolabs, Cat# M0202S) following the Golden Gate cloning method. The ligation product was isopopanol precipitated and electroporated into Stbl4 electrocompetent cells (Invitrogen, Cat# C751003) and grown at 30 °C. Plasmid DNA (pDNA) was prepared (HiSpeed Plasmid Maxi Kit, Qiagen, Cat# 12662) and pooled. Library representation and distribution were confirmed by next-generation sequencing.

#### Mutant library cell preparation

Lentivirus for pooled library production was generated by transfecting HEK293T cells (20×1e6) in a 225 cm^2^ flask with a DNA mixture containing pMD2.G (5 μg), psPAX2 (50 μg), and 40 μg of the library vector, using PEI (1 mg/mL) at a 1:3 ratio. The viral supernatant was collected 48 h post-transfection. To determine lentiviral titer, eHAP1-7TS cells were transduced with varying volumes of lentivirus. After 48 h, puromycin (5 µg/mL; Sigma-Aldrich, Cat# P8833) was added, and after five days, cell viability was assessed to determine the viral dose that resulted in a transduction efficiency of 30-50%, corresponding to a multiplicity of infection (MOI) of approximately 0.35-0.70. To generate mutant library cells, eHAP1-7TS cells were transduced in two biological replicates at a low MOI of approximately 0.5. Briefly, cells were seeded at a density of 1.5×10^6^ cells per well in a 12-well plate containing IMDM. 24 h post seeding, plates were subjected to spinfection with lentivirus mixed with polybrene (8 µg/mL) at 640xg for 2 h to enhance transduction efficiency, followed by a 4-6 h incubation after which cells were trypsinized and seeded into 225-cm^2^ flasks. Two days post-transduction, puromycin (5 μg/mL) was added and maintained for seven days to eliminate non-transduced cells.

#### FACS-based phenotype enrichment for pooled screens

For each biological replicate screen, mutant library cells were thawed in 15-cm dishes, split and seeded for the FACS sorting stage. Cells were treated with 50% Wnt-3a conditioned medium for 24 h and then sorted by flow cytometry to isolate the top 10% and bottom 15% of the reporter population, with unsorted cells collected as a reference for log-fold change normalization.

#### Next Generation Sequencing

Genomic DNA (gDNA) was isolated from sorted and unsorted cell populations using the Mag-Bind Blood & Tissue DNA HDQ Kit (Omega Bio-Tek, Cat# M6393). For PCR amplification of the integrated sgRNA sequences, 10 μg of gDNA was combined with a 100 µL reaction mixture containing 2x NEBNext Master Mix and primers containing adaptor sequences with unique barcoded i7 and i5 indices for Illumina sequencing. PCR cycling conditions were: (1) Initial denaturation: 95 °C for 1 min; (2) Denaturation: 94 °C for 30 s; (3) Annealing: 52.5 °C for 30 s; (4) Extension: 72 °C for 30 s; (5) cycle through steps 2-4 for 25 cycles; (6) Final extension: 72 °C for 5 min. PCR products were resolved on a 2% TAE gel, purified and library concentrations were estimated using qRT-PCR with PhiX (Illumina, Cat# FC-110-3001) as a standard. The samples were then sequenced on an Illumina NextSeq 550 platform with a 20% spike-in of PhiX.

#### Data analyses

Data analysis was performed using R (v4.3.2). Raw sequencing reads were demultiplexed and converted to FASTQ files using the bcl2fastq module (v2.20.0), with each file assigned to a condition based on the 8-nt P5 and P7 primers. These FASTQ files were then preprocessed to extract the 20-nt guide RNA sequences using the PreprocessReads function from the QuasR package (v1.36.0)^78^. The guide RNA reads were subsequently aligned to a custom reference file containing all possible guide RNA sequences in the library using the Subread algorithm from the Rsubread package (v2.16.2)^79^. Following alignment, a read count matrix was generated^80^. The raw read counts were normalized to reads per million (RPM) for each condition using the formula: RPM=(reads per guide RNA / total reads per condition)×1e6. To prevent issues with zero counts, a log_2_ transformation was applied to the RPM values after adding a pseudocount of 1 to all counts. We filtered out any sgRNAs with log₂-normalized reads per million in the unsorted sample falling outside 1.5 times the interquartile range (IQR) from the first and third quartiles, effectively removing outliers and guides with extremely low or high representation. The log-fold change (LFC) between sorted and unsorted populations was then calculated for each sgRNA. To provide a standardized score (z-score) of enrichment or depletion, we used the LFC values from control sgRNAs (non-targeting) to compute a mean and standard deviation, which were then used to calculate z-scores for each sgRNA in the respective sorted populations as described previously^27^. Predicted base editing mutations (N3-N10 window) were categorized using custom R functions. Each mutation was assigned a category—including “missense,” “nonsense,” “splice-site,” “silent,”—by parsing annotation strings derived from the library design. These categories were stored in a new column within the dataset for subsequent analysis and plotting. All analyses were conducted in R, and the complete code and analysis pipelines are available on GitHub to ensure reproducibility and transparency.

### Validation of base editor gRNA hits

To validate the top hits from the base editing screens, sgRNAs with z-scores ≥2 were selected. Individual targeting sequences were cloned into either the pRDA 429 or pRDA 336 plasmids. Screen hits were validated in two independent cell lines, eHAP1-7TS and HEK293T-7TS, with reporter line generation and culture conditions described in the preceding sections.

#### Validation in eHAP1-7TS cells

Lentivirus was produced as described above. For each sgRNA, 1.5×1e6 eHAP1-7TS reporter cells were seeded per well in a 12-well plate. 24 h post cell seeding, cells were transduced with 1.5 mL of lentivirus in a growth medium containing polybrene (8 μg/mL). The plates were centrifuged at 640×g for 2 h before being transferred to a 37 °C incubator. After 4–6 h, cells were trypsinized and transferred to a 10-cm dish. Puromycin selection was initiated 48 h post-transduction by replacing the growth medium with fresh medium containing puromycin (5 μg/mL) and continued for 5–7 days. Following selection, cells were split into two conditions: one in growth medium without WNT conditioned media and a second supplemented with 50% WNT conditioned media. After 24 h of treatment, cells were trypsinized, suspended in growth medium, and mScarlet fluorescence was measured by flow cytometry using a SH800S cell sorter.

#### Validation in HEK293T-7TS cells

sgRNAs that resulted in ≥50% reduction in reporter fluorescence (for hits from the bottom 15% sorted gate) and ≥10% elevation in reporter fluorescence (for hits from the top 10% sorted gate) were selected for an independent round of validation in HEK293T-7TS reporter cells. Cells were seeded at 3×10^5^ cells per well in a 12-well plate. 24 h post-seeding, cells were transfected with 1 μg of the pRDA plasmid using the PEI transfection reagent. 48 h after transfection, cells were trypsinized and seeded into 6-cm dishes. Puromycin selection was started 24 h later at 5 μg/mL and continued for 5–6 days. Cells were then subjected to WNT ligand treatment for 24 h and mScarlet fluorescence was measured as described for the eHAP1-7TS cells.

#### Data analysis and hit selection

Due to the large scale of the initial validation screen (370 sgRNAs), measurements were conducted as a single experiment in both eHAP1-7TS and HEK293T-7TS reporter cell lines. The selection of sgRNAs for subsequent, more detailed characterization was based on the criterion that a given sgRNA showed the same phenotype (either a reduction or an elevation in fluorescence) in both cell lines. sgRNAs that met this criterion were then subjected to detailed characterization in at least three independent experiments.

### FACS-based phenotype enriched mutant cell generation

Following the selection of hits for detailed characterization, mutant cell populations were generated through FACS-based enrichment. The procedure was performed on HEK293T-7TS mutant cells as described in the previous section. Two distinct protocols were followed based on the observed phenotype:

For hits that resulted in a reduction of reporter fluorescence, cells were treated with 25% Wnt-3a conditioned medium and 10 ng/mL RSPO3 for 24 h. Cells with the lowest mScarlet fluorescence peak were then sorted using flow cytometry.

For hits that resulted in an elevation of reporter fluorescence, cells were treated with 5% Wnt-3a conditioned medium and 10 ng/mL RSPO3 for 24 h. Cells with the highest mScarlet fluorescence peak were then sorted.

In both cases, approximately 1×10^6^ cells were sorted, expanded, and frozen for future analysis. To determine the enriched mutational signature that resulted in the phenotype, deep sequencing of the target region was conducted. This was performed as described in the ‘Next Generation Sequencing’ section, with the key difference being a two-step PCR protocol. The first step used primers with appended Illumina adapters, and the second PCR step incorporated primers with adapters and barcodes for demultiplexing (Supplementary Table 3). Reads were analyzed with CRISPResso2^81^ to determine allele frequencies in the pooled sorted cells.

### Recombinant *Sp*Cas9 and sgRNA purification and Ribonucleoprotein (RNP) complex assembly

Recombinant *Streptococcus pyogenes* Cas9 (*Sp*Cas9) protein was purified using a protocol adapted from the previously described method^82^. The expression plasmid pMJ915 (Addgene, plasmid#69090), which encodes *Sp*Cas9 with an N-terminal His-MBP-TEV tag and a C-terminal 2xSV40NLS sequence, was transformed into Rosetta 2(DE3)pLysS Competent *E. coli* cells (Millipore-Sigma,Cat#71403). *E. coli* were cultured in terrific broth (Thermo Scientific, Cat#H26824.36) at 37 °C. Protein expression was induced with 0.25 mM IPTG at a cell density of OD_600_=0.6, followed by incubation at 18 °C for 16 h. Cell pellets were harvested by centrifugation and resuspended in lysis buffer (20 mM Tris pH 8.0, 500 mM NaCl, 1 mM TCEP, 5% glycerol, 5 mM imidazole, 0.5 mM PMSF, and 1x protease inhibitor cocktail (Millipore-Sigma, Cat#S8820). Resuspended cells were homogenized using an EmulsiFlex-C5 (Avestin) at 20,000 psi. Lysates were clarified by centrifugation at 50,000xg for 30 minutes and the supernatant was incubated with His-Tag Purification Resin (Roche, Cat#5893682001) for batch binding. The resin was washed with ten column volumes of wash buffer (lysis buffer excluding PMSF and protease inhibitor cocktail). Bound protein was eluted with an elution buffer (wash buffer supplemented with 250 mM imidazole). TEV protease (homemade) was added to the eluted protein, and the mixture was dialyzed overnight against Ion-Exchange Buffer A (20 mM HEPES pH 7.5, 150 mM KCl, 1 mM TCEP, and 5% glycerol) using a 10 MWCO dialysis cassette (Thermo Fisher Scientific, Cat#66453) to cleave the His-MBP tag. The cleaved sample was then applied to a HiTrap SP HP column (cytiva) equilibrated in 15% Ion-Exchange Buffer B (20 mM HEPES pH 7.5, 1 M KCl, 1 mM TCEP, and 5% glycerol). *Sp*Cas9 was eluted with a linear gradient from 15% to 50% Ion-Exchange Buffer B. Peak fractions were pooled, concentrated using an Amicon ultracentrifugal filter, 30 kDa MWCO (Millipore-Sigma, Cat#UFC9030), and subjected to size exclusion chromatography on a HiLoad 16/60 Superdex 200 prep grade column (cytiva) equilibrated in Size Exclusion Buffer (20 mM HEPES pH 7.5, 150 mM KCl, 1 mM TCEP, and 10% glycerol). Peak fractions containing purified *Sp*Cas9 were pooled, aliquoted, flash frozen in liquid nitrogen, and stored at −80 °C. sgRNA template assembly, amplification, *in vitro* transcription, and purification were performed as described previously^82^. Target sequences that are part of the T7FwdVar oligo in the sgRNA template assembly step are listed in the “APCmcr truncation in HAP1 cells” and “PGE of AXIN1 in HAP1 cells” sections. *In vitro* transcription was carried out using the HiScribe T7 High yield RNA Synthesis Kit (New England Biolabs, Cat#E2040S) following the manufacturer’s instructions. Transcription reactions were subsequently treated with RNase-free DNase I (Thermo Fisher Scientific, Cat EN0521) to remove DNA template, followed by Quick CIP (New England Biolabs, Cat#M0525). sgRNAs were purified using the RNeasy Mini kit (Qiagen, Cat#74104) following a modified protocol^82^. For CRISPR/Cas9 mediated non-homologous end joining (NHEJ) and homology-directed repair (HDR) experiments, Cas9 ribonucleoprotein (RNP) complexes were assembled immediately before use. A 20 µL reaction volume was prepared containing 100 pmol of purified *Sp*Cas9 protein and 120 pmol of sgRNA in Cas9 buffer (20 mM HEPES pH 7.4, 150 mM KCl, 1 mM MgCl_2_, 1 mM TCEP, and 10% glycerol).

### APC^mcr^ truncation in RKO and HAP1 cells

Truncations within the mutation cluster region (MCR) of the endogenous APC gene were introduced into RKO and HAP1 cells using CRISPR/Cas9-mediated gene editing. RKO cells were prepared for electroporation using the Cell Line Nucleofector Kit R (Lonza Bioscience, Cat#VCA-1001). For electroporation reaction, 1 µg of the plasmid pSpCas9(BB)-2A-GFP (Addgene, Cat#48138), encoding *Sp*Cas9 and sgRNA targeting the APC^mcr^ (APC sg1320; 5’-ACTGCTGGAACTTCGCTCAC-3’), was added to the cell suspension. Electroporation was carried out using the Amaxa Nucleofector II system following the manufacturer’s instructions with program A-024. APC^mcr^ truncation mutation was generated in HAP1 cells by delivering pre-assembled Cas9 ribonucleoprotein (RNP) complexes via electroporation. The RNP complexes were assembled using purified *Sp*Cas9 protein and *in vitro* transcribed sgRNA targeting the APC^mcr^ (APC sg1336, 5’-ACCAAATCCAGCAGACTGCA-3’). HAP1 cells were electroporated with the assembled RNP complexes using the NEPA21 Electro-Kinetic transfection system (Bulldog-Bio). The specific electroporation parameters used were: poring pulse of 190 V, poring length of 5 ms, and the remaining parameters being default settings. Approximately 24 h after electroporation, the cell culture medium was replaced with fresh growth medium. On the following day (approximately 48 h post-electroporation), live single cells were sorted into individual wells of a 96-well plate containing growth medium using a SH800 Cell Sorter. Single-cell clones were subsequently expanded in culture. Successful introduction of APC^mcr^ truncation mutations in clonal cell lines was confirmed by both Sanger sequencing of the targeted APC locus and immunoblotting to assess the size of APC.

### Validation of base editor sgRNA hits in APC^mcr^ truncated cells

sgRNAs that passed the hit selection criteria from the “Validation of base editor sgRNA hits” section were cloned into either the pRDA 429 or pRDA 336 plasmids. Lentivirus production and transduction was performed as described in the ‘Generation of WNT transcriptional reporter lines’ section. The recipient clonal cell lines (RKO-7TG 1320* and DLD-1-7TG) were transduced at a multiplicity of infection (MOI) of approximately 0.5 in the presence of polybrene (8 µg/mL). Transduced cells were subjected to puromycin selection at a concentration of 5 µg/mL for 5-7 days, or until all non-transduced control cells had died.

### Precise genome editing (PGE) of *AXIN1* locus in HAP1 cells

sgRNAs and single-stranded oligonucleotide (ssODN) donor templates for precise genome editing of the *AXIN1* locus via homology directed repair (HDR) were designed using the CRISPR HDR Design Tool (Integrated DNA Technologies). The targeting sequences for *AXIN1* were 5’-GGAGAACCCTGAGAGCATCC-3’ (L471A/D472A/H474) and 5’-CAGCGTGTGCTGAGGACACC-3’ (L479W). ssODN donor templates were designed to introduce either the L471A/D472A/H474A or the L479W mutations into the *AXIN1* gene. The sequences of the ssODN templates are listed in the Supplementary Table 3. A silent mutation designed to introduce a *Pst*I restriction site was incorporated into the ssODN for the *AXIN1* L471A/D472A/H474A mutation to facilitate screening, designed using the silent mutagenesis tool (molbiotools). A mutation-specific forward primer was designed to screen for the *AXIN1* L479W mutant allele. HAP1 cells, both wild-type for APC (full length) and those harboring the APCmcr truncation, were subjected to electroporation to deliver the CRISPR/Cas9 RNP complex and ssODN donor templates for HDR. Electroporation was performed as described in the “generation of APCmcr truncation in HAP1 cells” section using the NEPA21 Electro-Kinetic transfection system. For each reaction, 100 pmol of pre-assembled *Sp*Cas9 RNP complex, 100 pmol of the respective ssODN donor template, and 1 µM of HDR Enhancer v2 (Integrated DNA Technologies, Cat#10007910) were added to the cell suspension immediately prior to electroporation. Single-cell sorting was performed approximately 72 h post-electroporation to isolate individual cells for clonal expansion, as described in the “single-cell sorting and clonal expansion” section. Sorting was performed based on viability gating. Putative positive clones were screened and confirmed for the presence of the desired *AXIN1* mutations. For clones putatively carrying the *AXIN1* L471A/D472A/H474A mutation, genomic DNA was extracted, and a region of the *AXIN1* locus was amplified by PCR. The PCR product was then digested with the *Pst*I restriction enzyme (New England Biolabs) to confirm the introduction of the silent mutation and thus the intended HDR event. Clones putatively carrying the *AXIN1* L479W mutation were screened by PCR using a mutation-specific forward primer (see Supplementary Table 3) followed by analysis of the PCR product size on an agarose gel. Positive clones identified by these screening methods were further confirmed by Sanger sequencing of the targeted *AXIN1* locus. To generate HAP1 cells harboring the *AXIN1* V475I/L479W double mutation, a clonal line previously established with the *AXIN1* L479W single mutation was used. This clonal line was transduced with a lentivirus generated from the plasmid pRDA 336 expressing a CBE targeting the *AXIN1* locus with the targeting sequence sequence 5’-CTGTACGTGCTCGTCCAGGA-3’. Following transduction, antibiotic selection was applied to enrich for transduced cells. Single-cell sorting, clonal expansion, and positive clone identification were performed as described earlier in the “single-cell sorting and clonal expansion” and “genotype confirmation” sections, respectively.

### Generation of doxycycline-inducible FLAG-AXIN1 stable SW480 cell lines

These cell lines were generated using the PiggyBac transposon system. Briefly, FLAG-AXIN1 wild-type, Affinity-Reducing (AR) mutant (L471A/D472A/H474A), and Affinity-Enhancing (AE) mutant (V475I/L479W) constructs were cloned into the PB-TAC-ERP2 (Addgene Cat#80478) plasmid. For transfection, 300,000 SW480 cells were seeded per well in a 6-well plate. Approximately 24 h later, the cells were transfected with 1 µg of the PB-TAC-ERP2 plasmid and 1 µg of the Super PiggyBac Transposase expressing plasmid using X-tremeGENE DNA 9 transfection reagent (Millipore-Sigma Cat#6365787001). At 48 h post-transfection, cells were trypsinized and re-plated for puromycin selection. Selection was performed with a concentration of 5 µg/mL of puromycin for 5 days.

### Generation of FLAG-AXIN1 stable SW480 cell lines by lentiviral transduction

FLAG-AXIN1 wild-type, Affinity-Reducing (AR) mutant (L471A/D472A/H474A), and Affinity-Enhancing (AE) mutant (V475I/L479W) constructs were cloned into the pRDA 336 plasmid by swapping the Cas9-NG sequence to retain the P2A-puromycin selection cassette. Lentivirus production and transduction was performed as described in the ‘Generation of WNT transcriptional reporter lines’ section. Recipient clonal cell lines were transduced at a multiplicity of infection (MOI) of approximately 0.5 in the presence of polybrene (8 µg/mL). Transduced cells were subjected to puromycin selection at a concentration of 5 µg/mL for 5 days.

### Immunoblotting

All cells were harvested by scraping. Samples were lysed according to the target protein to ensure optimal extraction. Protein concentrations were determined using the BCA Protein Assay Kit (Thermo Fisher Scientific, Cat#23225) with bovine serum albumin (BSA) as the standard. For the analysis of soluble (non-phosphorylated, active) β-catenin, cells were lysed in hypotonic buffer (10 mM Tris pH 7.5, 10 mM KCl, 5 mM EDTA) supplemented with a Protease and Phosphatase Inhibitor cocktail (Thermo Fisher Scientific, Cat#78442) and 1 μM Bortezomib. After a 10-minute incubation on ice, cells were lysed via a freeze-thaw cycle using liquid nitrogen. Lysates were then centrifuged at 20,000 x *g* for 30 minutes at 4 °C to separate the soluble fraction, and the supernatant was collected for analysis. For the analysis of AXIN1, cells were lysed in a Cell Lysis Buffer with Ionic and Non-Ionic Detergents (CLB-wIND; 50 mM Tris pH 8.0, 150 mM NaCl, 1% Triton X-100, 0.1% SDS, 0.1% Deoxycholate, 10% glycerol) containing a Protease and Phosphatase Inhibitor cocktail. Samples were vortexed for 30 minutes at 4 °C to ensure complete lysis. Lysates were then clarified by centrifugation at 20,000 x *g* for 30 minutes at 4 °C, and the resulting supernatant was collected for analysis. For immunoblotting, 20–30 μg of total protein was separated by SDS-PAGE using 4–12% gradient NuPAGE Bis-Tris Protein Gels (Thermo Fisher Scientific). Separated proteins were transferred to a nitrocellulose membrane (Bio-Rad, Cat#1620112). Membranes were blocked for 30 minutes at room temperature in either LI-COR Odyssey Blocking Buffer (0.2x PBS, 0.1% casein, and 0.01% sodium azide) for β-catenin blots or Blocking Buffer M (Tris-buffered saline (TBS) containing 0.1% Tween-20 (TBS-T) and 5% non-fat dry milk) for AXIN1 blots. Primary antibodies were incubated overnight at 4 °C. After three washes with TBS-T, membranes were incubated for 1 h at room temperature with the appropriate secondary antibody: IRDye 800CW (LICORbio) for β-catenin or HRP-conjugated (Jackson ImmunoResearch) for AXIN1. Blots were washed three times with TBS-T. β-catenin blots were imaged using an Odyssey CLx Imaging System (LICORbio). Chemiluminescence for AXIN1 was detected using SuperSignal West Pico PLUS chemiluminescent substrate (Thermo Fisher Scientific, Cat#34580), and signals were captured on film.

### Co-immunoprecipitation

Cells were harvested by scraping and lysed in a Cell Lysis Buffer with Non-Ionic Detergent (CLB-wND; 50 mM Tris pH 7.4, 150 mM NaCl, 1% Triton X-100, 1 mM DTT, 10% glycerol) supplemented with a Protease and Phosphatase Inhibitor cocktail (Thermo Fisher Scientific, Cat#78442), 1 μM Bortezomib, 2 mM MgCl_2_, and 50 U/mL Benzonase. Lysates were rotated on a nutator for 1 h at 4 °C and then clarified by centrifugation at 20,000 x *g* for 30 minutes at 4 °C. For the immunoprecipitation of endogenous proteins (β-catenin and AXIN1), 3–4 mg of clarified cell lysate was pre-cleared with Dynabeads Protein A/G (Thermo Fisher Scientific, Cat#10002D and Cat#10004D) for 1 h at 4 °C to remove proteins that non-specifically bind to the beads. Following this, the pre-cleared lysates were incubated overnight at 4 °C on a nutator with pre-conjugated antibody-bead complexes. For β-catenin immunoprecipitation, 2 μg of β-catenin antibody was conjugated to Dynabeads Protein G. For AXIN1 immunoprecipitation, 2 μg of AXIN1 antibody was conjugated to Dynabeads Protein A. The resulting immunocomplexes were captured and washed three times with wash buffer (CLB-wND without inhibitors). Immunocomplexes were then eluted in 2x Laemmli sample buffer. For the immunoprecipitation of FLAG-tagged AXIN1 in SW480 cells, the procedure was identical, with the exception that anti-FLAG M2 magnetic beads (Sigma-Millipore, Cat#M8823) were used instead of the antibody-bead conjugation step and the pre-clearing step was omitted due to the low non-specific binding of the beads. Eluted proteins from all immunoprecipitation experiments were analyzed by immunoblotting.

### Immunofluorescence

Cells cultured on glass coverslips were fixed with 4% paraformaldehyde (PFA) in PBS for 10 min at room temperature (r/t). Following fixation, cells were permeabilized and blocked for 30 minutes at r/t using Blocking and Permeabilization Buffer (BPB) (1x PBS pH 7.4 containing 1% donkey serum, 1% BSA, and 0.1% Triton X-100). Coverslips were then incubated with the primary antibody against active β-Catenin (diluted in BPB) for 1 h at r/t. After three washes with PBS, cells were incubated with Alexa Fluor 488 conjugated donkey anti-rabbit IgG secondary antibody (Thermo Fisher Scientific) for 45 minutes at r/t in the dark. Coverslips were washed three times with PBS and mounted onto glass slides using ProLong Diamond Antifade Mountant with DAPI (Invitrogen, Cat#P36962) to counterstain nuclei. Images were acquired on a Leica TCS SP8 confocal microscope using a 63× oil-immersion objective. Z-stacks covering the entire cell were collected at 0.5 µm intervals, and maximum intensity projection images were generated for analysis. Nuclear and cytoplasmic fluorescence intensities were quantified using CellProfiler (v4)^83^. Nuclear masks were defined by automated thresholding of the DAPI channel, and cytoplasmic masks were generated by radial expansion of nuclear boundaries while excluding neighboring nuclei. Mean fluorescence intensities were extracted from nuclear and cytoplasmic compartments for downstream analyses.

### Quantitative real-time PCR

Total RNA was extracted from cells using TRIzol reagent (Thermo Fisher Scientific, Cat#15596026) following the manufacturer’s protocol. The RNA concentration and purity were determined using a Synergy H1 microplate reader (Biotek). cDNA was synthesized from 250 ng of total RNA using the iScript Reverse Transcription Supermix (Bio-Rad, Cat#1708841). Quantitative real-time PCR was performed on a QuantStudio 5 Real-Time System (Agilent) using SYBR Green qPCR Master Mix (Selleck, Cat#B21703) following the manufacturer’s instructions. A melt curve analysis was performed at the end of the run to ensure product specificity. The expression of target genes was normalized to the housekeeping gene *GAPDH*. The primers used were: hAXIN2_Fwd: 5’ – GTCCAGCAAAACTCTGAGGG – 3’, hAXIN2_Rev: 5’ – CTGGTGCAAAGACATAGCCA – 3’, hGAPDH_Fwd: 5’ – TGCACCACCAACTGCTTAGC – 3’, hGAPDH_Rev: 5’ – GGCATGGACTGTGGTCATGAG – 3’.

### Expression and purification of recombinant proteins

Recombinant proteins were expressed in *E. coli* using the same protocol as described for the *Sp*Cas9 protein in the ‘Recombinant *Sp*Cas9 and sgRNA purification’ section. For the GST-tagged β-catenin binders (AXIN1, a.a. Cys435-His498; TCF4, a.a. Met1-Glu53; ICAT, a.a. Met1-Gln81, cell pellets were resuspended in B-PER Reagent (Thermo Fisher Scientific, Cat#90079) supplemented with lysozyme (100 µg/mL), DNase I (5 U/mL), 1 mM PMSF, and protease inhibitor cocktail (Sigma-Millipore) and incubated for 15 min at room temperature. The lysate was clarified by centrifugation at 50,000 x *g* for 30 minutes at 4 °C. For affinity chromatography, clarified lysates were incubated with Glutathione Sepharose resin (Gold Biotech) for 1 h at 4 °C on a rotator. The mixture was transferred to a gravity-flow chromatography column (Bio-Rad), and the column was washed with 50 column volumes of GST-Wash Buffer (50 mM sodium phosphate buffer pH 7.4, 500 mM NaCl and 1 mM DTT). The GST-tagged proteins were then eluted with GST-Elution Buffer (50 mM sodium phosphate buffer pH 7.4, 150 mM NaCl, 1 mM DTT and 10 mM L-Glutathione). The eluted proteins were buffer exchanged into GST-Storage Buffer (30 mM sodium phosphate buffer pH 7.4, 150 mM NaCl, 1 mM DTT, 5% glycerol) using an Amicon Ultra-15 Centrifugal Filter Unit (Millipore, Cat#UFC901024, 10 kDa MWCO). Protein concentration was determined by measuring the absorbance at 280 nm. To purify GST-brSUMO-tagged full-length β-catenin, cell pellets were resuspended in GST-Lysis Buffer (50 mM Sodium phosphate buffer pH 7.4, 150 mM NaCl, 1 mM DTT) supplemented with 1 mM PMSF, and protease inhibitor cocktail. The cells were lysed by high-pressure homogenization (LM20 Microfluidizer) with two passes at 20,000 psi. The lysate was clarified by centrifugation at 50,000 x *g* for 30 minutes at 4 °C. The clarified lysate was incubated with Glutathione resin for 1 h at 4 °C on a rotator and then transferred to a gravity-flow column. The column was washed with 50 column volumes of GST-Wash Buffer, followed by a wash with 10 column volumes of Cleavage Buffer (50 mM Sodium phosphate buffer pH 7.4, 150 mM NaCl and 1 mM DTT). GST-brSUMO tag was cleaved by incubating the sample with 200 nM bdSENP1 (homemade) for 1 h at 4 °C on a rotator. The cleaved protein was then eluted using a gravity-flow column, and the eluate was centrifuged at 20,000 x *g* for 10 minutes at 4 °C. The sample was subjected to a size-exclusion chromatography on a HiLoad 16/60 Superdex 200 prep grade column (cytiva) equilibrated in SEC Buffer (30 mM Sodium phosphate buffer pH 7.4, 150 mM NaCl, 5% glycerol). Fractions containing the purified protein were pooled and concentrated using an Amicon 30 kDa MWCO concentrator. Protein concentration was estimated by measuring the absorbance at 280 nm.

### APC-AXIN1 pull-down assays to explore the contribution of β-catenin

Human AXIN1, β-catenin and APC variants were expressed using the baculovirus expression system in Sf9 cells and purified as described previously^16^. For the *in vitro* binding (pull-down) assay, preys (APC variants, β-catenin) were added to AXIN1-dStrepII (the bait) in equimolar ratios and left to incubate at room temperature for 20 min in binding buffer (50 mM HEPES pH 7.5, 150 mM NaCl, 5% glycerol, 1 mM TCEP). Then, 30 µl of packed StrepTactin Sepharose resin (IBA Lifesciences) was added to the binding reactions in the binding buffer to a total volume of 500 µl and left to incubate for 60 min at 4°C rotating. The resin was washed three times with 1 ml of binding buffer, allowing 5 min for incubation at each wash step. Finally, 50 µl of 1.5x SDS sample buffer was added to the resin. Samples were boiled and analyzed by SDS-PAGE and Coomassie staining.

### Pull-down assays

To investigate the *in vitro* binding interaction between GST-AXIN1 CBD and full-length β-catenin, we performed pull-down assays. *In vitro* binding reactions were set up in a total volume of 100 µL using a binding buffer containing 20 mM Tris pH 7.5, 150 mM NaCl, 0.1% Triton X-100, and 2 mM DTT. A fixed concentration of 1 µM GST-AXIN1 CBD was incubated with increasing concentrations of full-length β-catenin (0.01, 0.1, 0.5, and 2 μM) for 30 min at 22 °C. Following the initial binding reaction, Glutathione Magnetic Agarose beads (Thermo Fisher Scientific, Cat# 78601) were added to each sample and incubated for an additional 30 min at 22 °C. The beads were then washed three times with a binding buffer to remove unbound proteins. The protein complexes were eluted by boiling in a 2x Laemmli sample buffer. Samples, representing 10% of the total elution volume, were analyzed by SDS-PAGE followed by immunoblotting with antibodies against GST and β-catenin, as described in the ‘Immunoblotting’ section.

### Biolayer interferometry-based protein binding assays

Kinetic binding experiments were performed on an Octet RED96 system (ForteBio, Sartorius) at 30 °C with a plate shaking speed of 1,000 rpm in 96-well black, flat-bottom polypropylene microplates (Greiner Bio-One, #655209). All assays were conducted in Octet buffer composed of 30 mM sodium phosphate pH 7.4, 150 mM NaCl, 1 mM DTT, and 5% glycerol, supplemented with 0.05% Tween-20 and 0.1% BSA to minimize nonspecific binding. Octet GST biosensor tips were pre-equilibrated in Octet buffer for 10 min prior to use. Sensors were then loaded with GST-tagged ligands (GST-AXIN1 and its variants, GST-ICAT, and GST-TCF4) at 50 nM, ensuring measurements remained within the linear range of the BLI signal to avoid overcrowding on the sensor surface. Following ligand immobilization, sensor tips were washed and baselined in wells containing only Octet buffer. For analyte binding, full-length β-catenin and its variants were used at concentrations of 500 nM for GST-ICAT, 1 µM for GST-Axin1 and GST-TCF4. Association was monitored for 20–25 s, followed by 100 s of dissociation in the Octet buffer. A reference sensor (ligand-loaded but dipped into buffer only) was included to correct for nonspecific signal drift and baseline fluctuations. All sensorgrams were reference-subtracted and analyzed using ForteBio Data Analysis Software 10.0 software. Experimental data from both association and dissociation phases was simultaneously fit using a 1:1 binding model to estimate *k*_on_ and *k*_off_. The association phase is modelled using pseudo-first-order kinetics, assuming that the analyte concentration is effectively constant since we used a ∼10–60 fold excess of analyte over immobilized ligand. The dissociation phase was modeled using first-order decay kinetics. The equilibrium dissociation constant was calculated as *K*_D_=*k*_off_/*k*_on_. Reported *K*_D_ values represent the average of at least three independent experiments.

### Data visualization, statistics and reproducibility

Quantitative data were visualized using GraphPad Prism and R. All structural figures were generated using UCSF ChimeraX (version 1.8rc202405140121). PDB codes for structural figures are provided in the respective figure legends. Immunoblot panels were processed using Adobe PhotoShop (for chemiluminescence data) and Fiji (for LI-COR data). Figure panels were prepared using Adobe Illustrator. Unless otherwise indicated, all experiments were performed with a minimum of three (n≥3) independent biological replicates. Technical replicates were not counted as independent samples for statistical analysis. Data are consistently presented as the mean with individual data points overlaid. For experiments involving comparisons between two groups (e.g., wild-type vs. a single mutant β-catenin variant) within a single treatment condition (untreated, Wnt-3a, or CHIR99021), statistical significance was determined using multiple unpaired Student’s *t*-tests with Welch’s correction. The rejection thresholds for the *p*-values were adjusted using the two-stage step-up (Benjamini, Krieger, and Yekutieli) method to control the false discovery rate (FDR) at 0.01. For experiments involving comparisons across multiple groups (e.g., wild-type β-catenin and four mutant variants) within a single treatment condition (untreated, Wnt-3a, or CHIR99021), statistical significance was determined using Brown-Forsythe and Welch Analysis of Variance (ANOVA) tests. Subsequent pairwise comparisons between groups were conducted using Dunnett’s T3 multiple comparisons test. Specific *p*-values for key comparisons are reported directly in the figure panels.

### Use of Artificial Intelligence

The authors used AI assistants to strategically refine different sections of this manuscript. The abstract was improved with the assistance of Claude (Anthropic) to enhance its appeal to a broader audience and convey the work’s significance beyond a specialized field. Gemini (Google) was used to ensure the methods and figure legends were detailed, clear, and consistent to promote greater reproducibility. The AI assistants did not perform any scientific analysis, interpretation of results, or contribute to the intellectual content of this research. All information presented is the sole product of the authors’ work and judgment.

## Data availability

All raw and processed sequencing data supporting the CRISPR-based screening and subsequent analyses are available in the Sequence Read Archive (SRA) repository under BioProject accession number PRJNA1344315. The specific read counts and processed data files derived from the initial base editing screen are listed in Supplementary Table 1. All source data and corresponding raw quantification files used for the validation of sgRNAs in eHAP1-7TS and HEK293T cell lines are provided in Supplementary Table 2, accompanying the paper.

## Code availability

The custom R scripts and computational pipelines utilized for the processing and statistical analysis of the base editing screen data were written in R (4.3.2) and are publicly available at the following GitHub repository:https://github.com/RohatgiLab/Murugesh_DC_base_editing_analysis

## Acknowledgements

We are grateful to Yuki Yoshida for cloning the GST-TCF4 CBD construct and to Ramin Dubey for generating the DLD-1-7TG WNT transcriptional reporter line. This work was supported by the National Institutes of Health (NIH) grants GM118082 (to R.R.), CA230220 (to S.M.C.), and GM122516, CA281002, and CA236733 (to E.L.). R.R. and E.L. were also supported by an Innovation Fund Award from the Pew Charitable Trusts. M.M. was supported by an American Cancer Society – Jean Perkins Foundation Postdoctoral Fellowship in Cancer Research (Award PF-20-121-01-TBE). Work in S.G. ‘s group was funded by a Cancer Research UK Programme Foundation Award (C47521/A28286) and a PhD Studentship by The Institute of Cancer Research to S.S.

## Author contributions

R.R. and G.V.P. conceived the study. M.P. optimized the base editing methodology, conducted screens, performed formal analyses, and generated the structure panels. M.P. also performed all Biolayer Interferometry (BLI) experiments, generated the eHAP1-7TS reporter line, and performed validation experiments in eHAP1-7TS cells. G.V.P. designed the experiments, engineered the HAP1 clonal lines using HDR, and performed mechanistic analyses of the engineered cell lines. G.V.P. also designed the sorting strategy for the HEK293T-7TS mutant cells and conducted all validation experiments in the HEK293T-7TS, RKO APC1320*-7TG, and DLD-1-7TG cell lines. M.P. performed the deep sequencing to determine allele frequency in the sorted HEK293T-7TS mutant cells. G.V.P. and R.R. purified recombinant proteins from bacteria. S.S. performed the *in vitro* AXIN1-APC-β-catenin pulldown experiments under the guidance of S.G.. P.S. and M.M. assisted with immunofluorescence staining and immunoblotting experiments. E.L. provided essential reagents. S.M.C. and M.P. contributed to the sgRNA library design. G.V.P. contributed to the data visualization with input from M.P. R.R. drafted the original manuscript with input from G.V.P. and M.P. R.R. and G.V.P. reviewed and edited the manuscript. R.R. supervised the study and acquired funding. All authors reviewed the manuscript.

## Competing interests

E.L. is a co-founder and holds equity in StemSynergy Therapeutics, a company developing inhibitors for major signaling pathways (including the WNT pathway) for the treatment of cancer. E.L.’s interests are reviewed and managed by Vanderbilt University in accordance with their conflict of interest policies. The remaining authors declare that they have no competing financial and non-financial interests.

**Correspondence and request for materials** should be addressed to Rajat Rohatgi.

---

**Extended Data Fig. 1.**
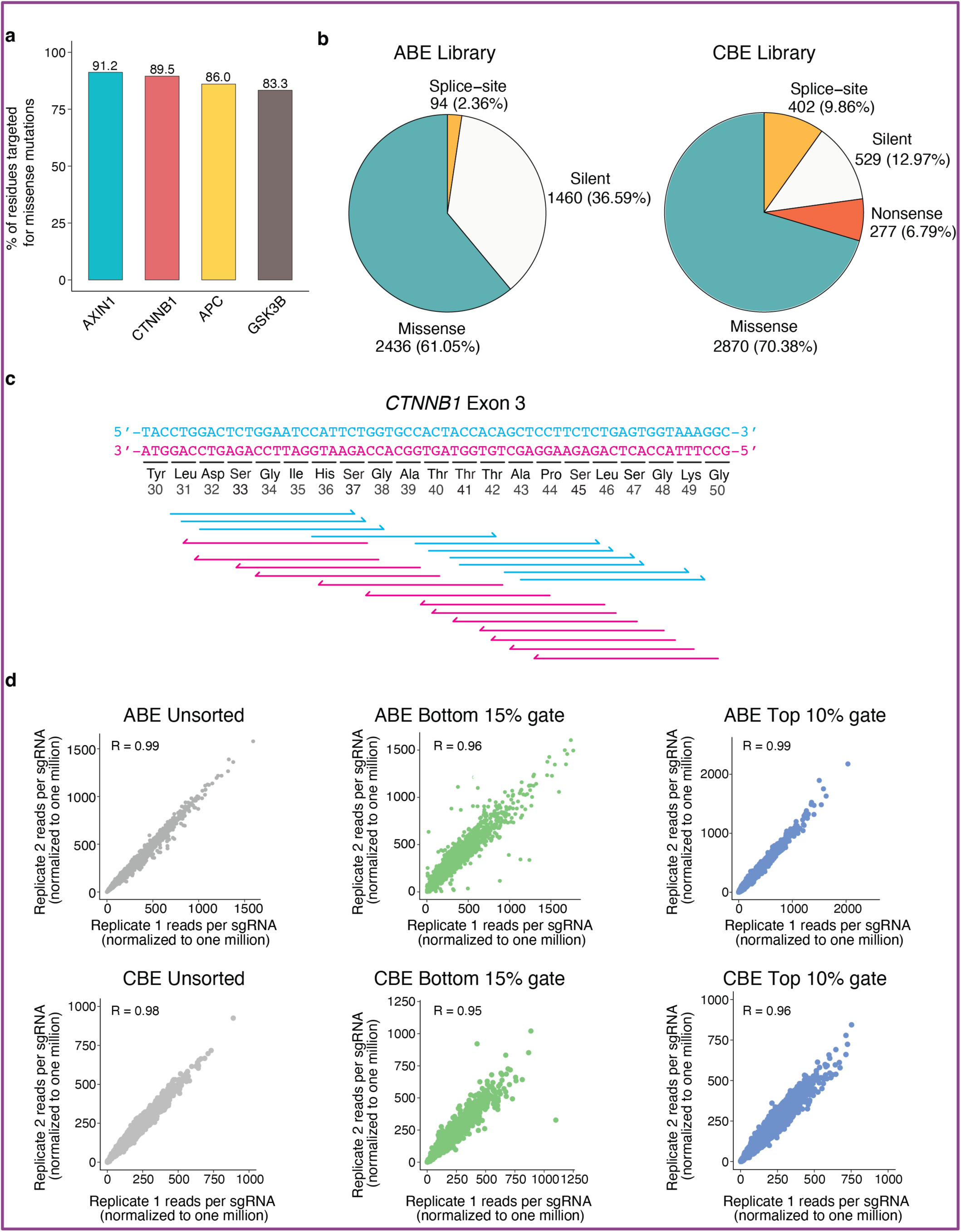
Design and quality control of ABE and CBE base editing screens targeting the β-catenin destruction complex (BDC). **a,** Bar graph summarizing the percent of amino acids in each BDC component predicted (i.e. located within the N3-N10 region of the protospacer) to be changed to a different amino acid by at least one sgRNA in our ABE and CBE libraries. **b,** Predicted proportion of missense, nonsense, silent, and splice-site mutations generated by the ABE (left) and CBE (right) libraries. Note that only CBE sgRNAs are predicted to generate nonsense mutations. **c,** The tiling nature of base editing libraries is illustrated using Exon 3 of *CTNNB1*. Arrows below the exon sequence show the position of sgRNA sequences, colored according to whether they hybridize to the top strand (red guides) or bottom strand (blue guides). Note that each codon in the exon is often targeted by multiple sgRNAs. **d,** Scatter plots showing the correlation of sgRNA read counts (per million total reads) between the two biological replicates for each of the six screens. See also Supplementary Table 1

**Extended Data Fig. 2.**
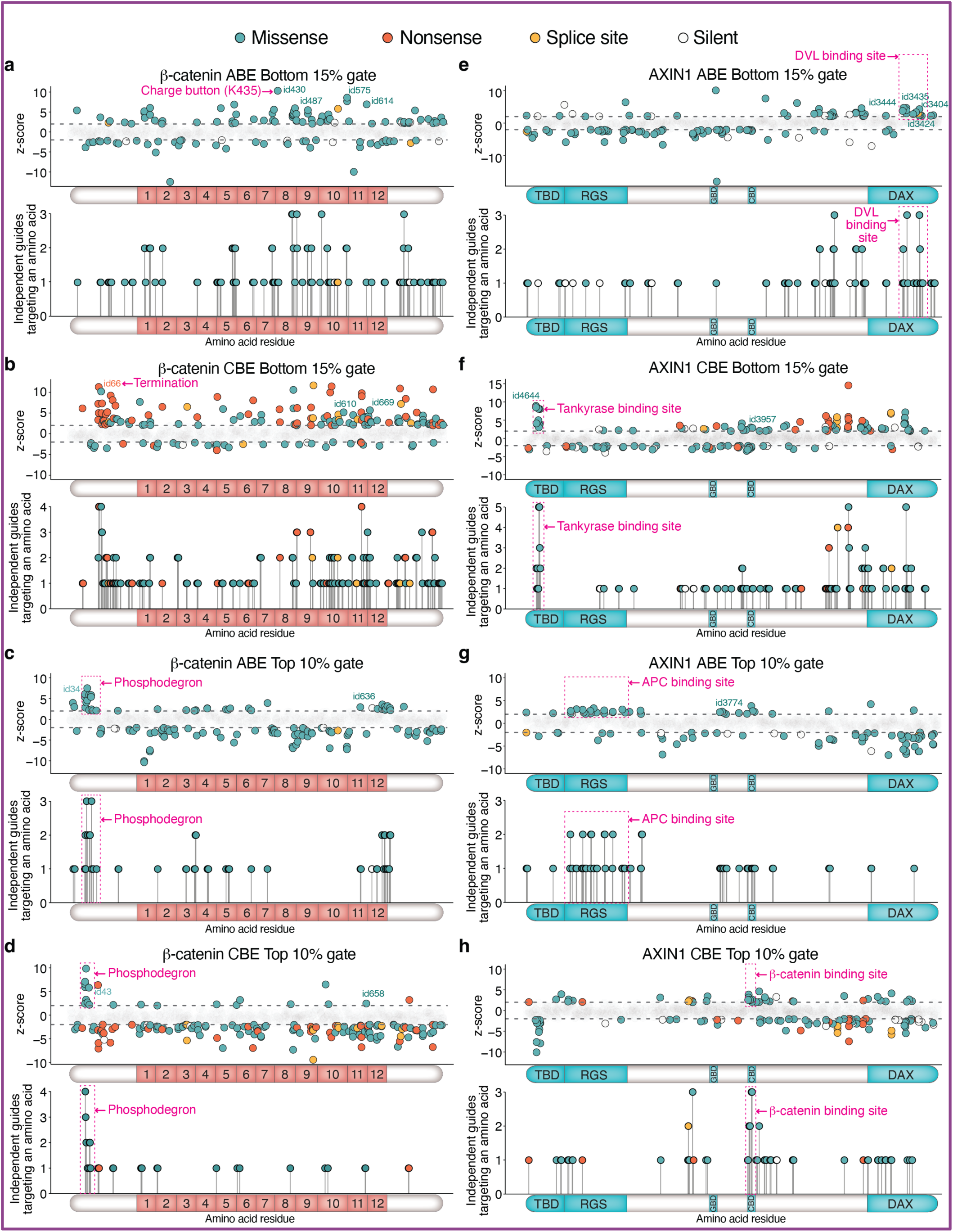
Sequence-function landscape of β-catenin and AXIN1 as mapped by tiled base editing screens. Each of the individual panels (**a-h**) show the position of all sgRNAs targeting the indicated protein using a z-score plot (top) and a lollipop plot (bottom). In the z-score plots, each sgRNA (represented by a point) is plotted based on its z-score for enrichment in the specified gate used for sorting (**Fig.1d**) on the y-axis and its amino acid position on the x-axis. For sgRNAs predicted to make multiple mutations, the most N-terminal residue is used as the position. A positive z-scores denotes enrichment in the indicated sorting gate; a negative z-score denotes depletion. sgRNAs with z-scores > 2 or < −2 (the criteria we used for statistical significance and follow-up) are shown as solid dots colored according to the type of predicted mutation (legend provided at the top of the figure). The target zone for each sgRNA was considered to span N3-N10 in its protospacer sequence. sgRNAs with −2< *z*-scores <2 are shown as faded dots and considered non-significant. In the lollipop plots, the y-axis represents the number of sgRNAs with z-scores ≥ 2 predicted to target a specific amino acid. In cases where an sgRNA was predicted to generate both missense and nonsense mutations, only the nonsense mutation is plotted. In all panels, magenta arrows and dotted squares point to known functional regions of the protein involved in key regulatory PTMs (e.g. the β-catenin phosphodegron) or PPIs. **a,b,** *z*-score and lollipop plots for β-catenin sgRNAs in the Bottom 15% gate (used to isolate mutant cells with reduced WNT/β-catenin signaling strength) from adenine (**a**, ABE) and cytosine (**b**, CBE) base editor screens. **c,d,** *z*-score and lollipop plots for β-catenin sgRNAs in the Top 10% gate (used to isolate mutant cells with elevated WNT/β-catenin signaling strength) from adenine (**c**, ABE) and cytosine (**d**, CBE) base editor screens. **e,f,** *z*-score and lollipop plots for AXIN1 sgRNAs in the Bottom 15% gate (used to isolate mutant cells with reduced WNT/β-catenin signaling strength) from adenine (**e**, ABE) and cytosine (**f**, CBE) base editor screens. **g,h,** *z*-score and lollipop plots for AXIN1 sgRNAs in the Top 10% gate (used to isolate mutant cells with elevated WNT/β-catenin signaling strength) from adenine (**g**, ABE) and cytosine (**h**, CBE) base editor screens. See also Supplementary Table 1

**Extended Data Fig. 3.**
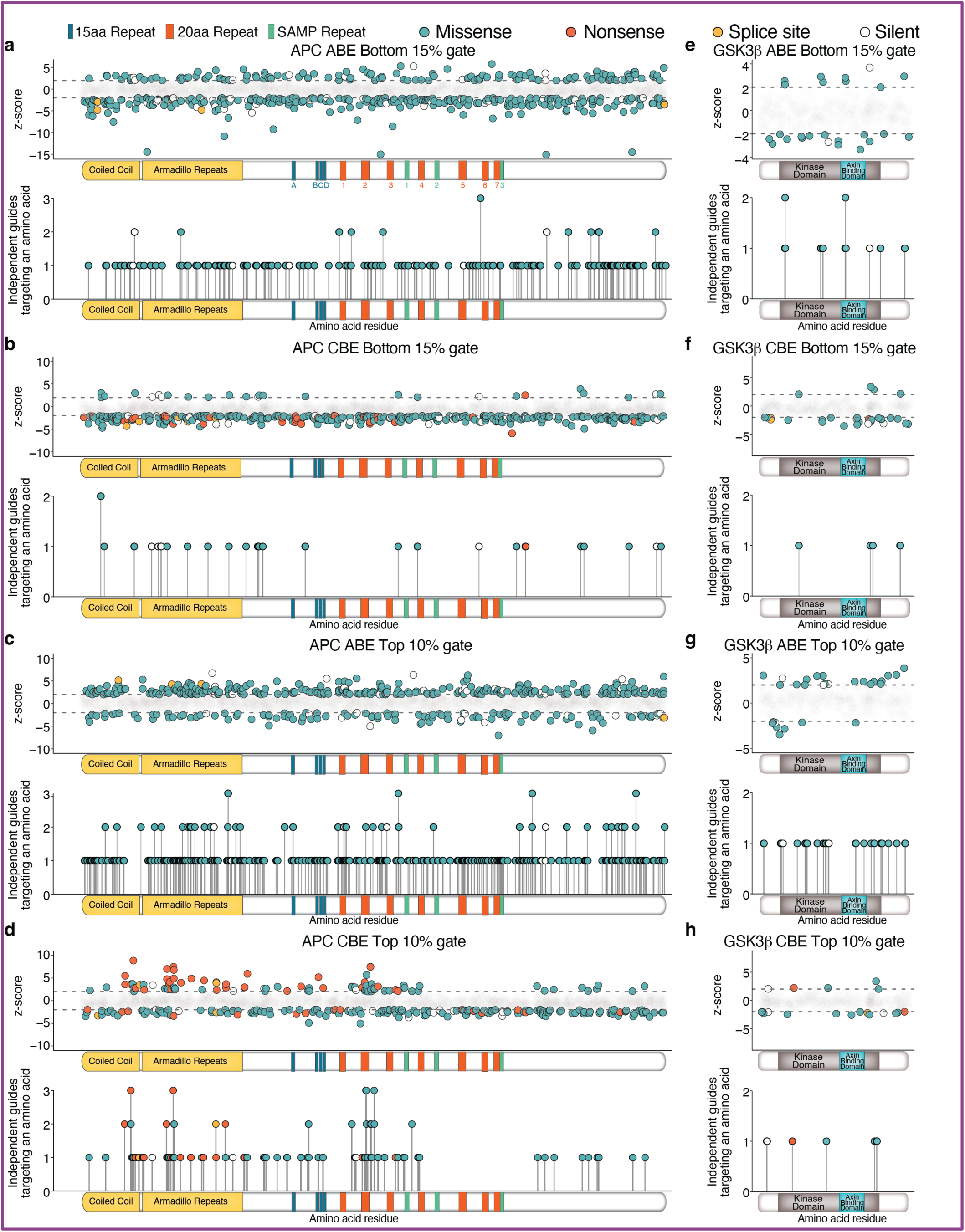
Sequence-function landscape of APC and GSK3β as mapped by tiled base editing screens. Each of the individual panels (**a-h**) show the position of all sgRNAs targeting the indicated protein using a z-score plot (top) and a lollipop plot (bottom). In the z-score plots, each sgRNA (represented by a point) is plotted based on its z-score for enrichment in the specified gate used for sorting (**Fig.1d**) on the y-axis and its amino acid position on the x-axis. A positive z-scores denotes enrichment in the indicated sorting gate; a negative z-score denotes depletion. sgRNAs with z-scores < 2 or < −2 (the criteria we used for statistical significance and follow-up) are shown as solid dots colored according to the type of predicted mutation (legend provided at the top of the figure). The target zone for each sgRNA was considered to span N3-N10 in its protospacer sequence. sgRNAs with −2< *z*-scores <2 are shown as faded dots and considered non-significant. In the lollipop plots, the y-axis represents the number of sgRNAs with z-scores ≥ 2 predicted to target a specific amino acid. In cases where an sgRNA was predicted to generate both missense and nonsense mutations, only the nonsense mutation is plotted. **a,b,** *z*-score and lollipop plots for APC sgRNAs in the Bottom 15% gate (used to isolate mutant cells with reduced WNT/β-catenin signaling strength) from adenine (**a**, ABE) and cytosine (**b**, CBE) base editor screens. **c,d,** *z*-score and lollipop plots for APC sgRNAs in the Top 10% gate (used to isolate mutant cells with elevated WNT/β-catenin signaling strength) from adenine (**c**, ABE) and cytosine (**d**, CBE) base editor screens. **e,f,** *z*-score and lollipop plots for GSK3β sgRNAs in the Bottom 15% gate (used to isolate mutant cells with reduced WNT/β-catenin signaling strength) from adenine (**e**, ABE) and cytosine (**f**, CBE) base editor screens. **g,h,** *z*-score and lollipop plots for GSK3β sgRNAs in the Top 10% gate (used to isolate mutant cells with elevated WNT/β-catenin signaling strength) from adenine (**g**, ABE) and cytosine (**h**, CBE) base editor screens. See also Supplementary Table 1

**Extended Data Fig. 4.**
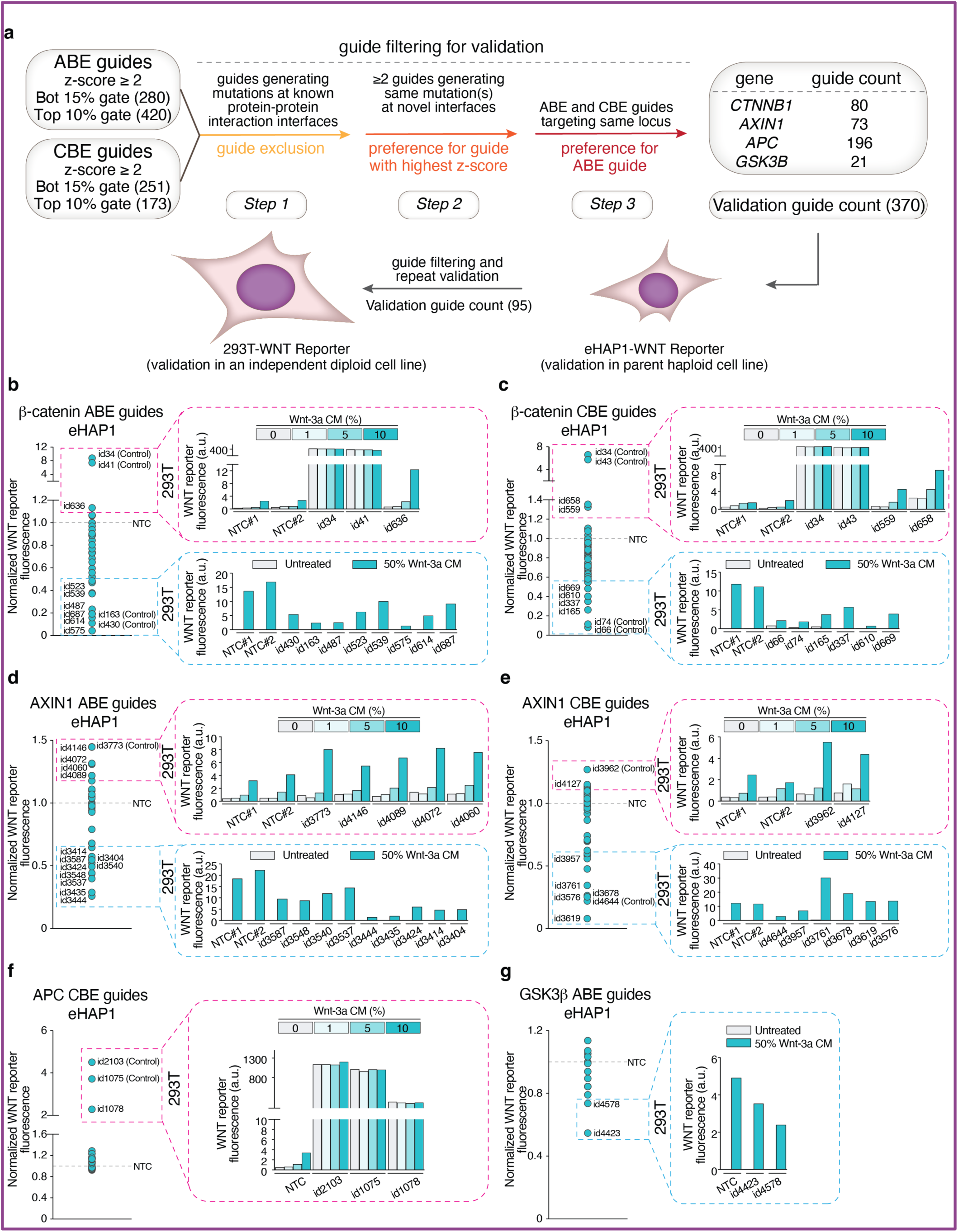
Validation strategy for hits from base editing screens. **a**, Outline of filtering and validation pipeline for sgRNAs with *z*-scores ≥ 2. After excluding most sgRNAs predicted to direct mutations at known PPI interfaces, remaining sgRNAs were individually introduced into eHAP1-7TS and HEK293T-7TS cells and WNT/β-catenin signaling strength assessed using the 7TS reporter. **b,c,** Validation of β-catenin sgRNAs from the ABE (**b**) and CBE (**c**) screens in both eHAP1-7TS and HEK293T-7TS reporter cells. sgRNAs were first tested for WNT/β-catenin signaling effects in eHAP1-7TS cells. Those that significantly increased (dotted magenta box) or decreased (dotted cyan box) reporter activity relative to non-targeting control sgRNA (NTC) were then retested in HEK293T-7TS cells (magenta and cyan insets). **d,e,** Validation of AXIN1 sgRNAs from the ABE (**d**) and CBE (**e**) screens, depicted like the β-catenin validation in **b** and **c**. **f,g,** Validation of *APC* sgRNAs from the CBE screen (**f**) and GSK3β sgRNAs from the ABE screen in both cell lines. eHAP1-7TS cells were treated with 25% Wnt-3a conditioned media (24h) before measuring 7TS reporter activity by flow cytometry. HEK293T-7TS cells carrying sgRNAs that decreased 7TS reporter activity were treated with 25% Wnt-3a conditioned media (24h). HEK293T-7TS cells carrying sgRNAs that increased 7TS reporter activity were treated (24h) with an increasing concentration of Wnt-3a conditioned medium (as shown in the magenta insets). All validated sgRNAs had directionally similar effects in both cell lines, except for AXIN1 CBE guides in panel **e** (ids: 3761, 3678, 3619, 3576), which were eliminated from consideration. See also Supplementary Table 2.

**Extended Data Fig. 5.**
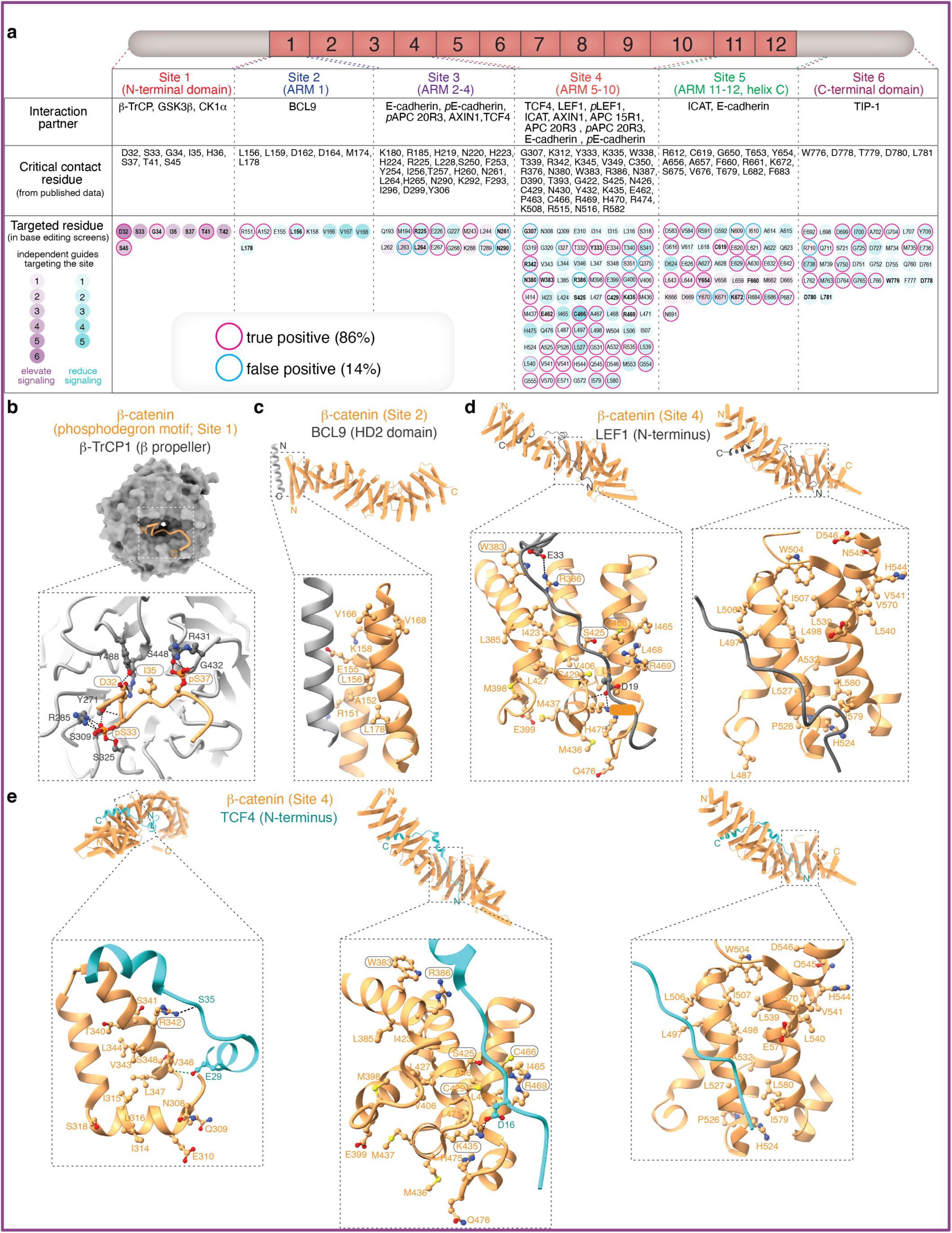
Validation of sgRNAs targeting β-catenin. **a,** Linear cartoon of β-catenin showing its 12 ARM repeats that form an extended ARM superhelix (see **Fig.2a**). An aligned table below shows the boundaries of Sites 1-6, known interaction partners at each Site (Row 1), β-catenin amino acids residues identified as playing a functional role at PPI interfaces in prior structural or biochemical studies (Row 2), and amino acid residues located in the N3-N10 segment of the protospacers of sgRNAs with a *z*-score ≥ 2 in our base editing screens (Row 3). Screen-identified residues are encircled by a dot colored teal for guides that reduce signaling and purple for guides that elevate signaling. The purple or teal shading intensity reflects the number of different sgRNAs predicted to target that amino acid residue (see scale in left-most column). Color of the ring surrounding each dot denotes whether it was a true positive (magenta) or false positive (blue) based on validation studies (**Extended Data Fig.4**); dots without a surrounding ring were not validated. Bold lettering denotes amino-acid residues that have been implicated in β-catenin function in a prior publication (e.g. the charge button K435). **b**, Structural model of the complex between the β-catenin phosphodegron motif (PDB 2Z6H, sandy brown) and the beta-propeller domain of β-TrCP1 (PDB 1P22, dark grey)^44^. In the zoomed view, side chains of β-catenin residues targeted by sgRNAs that elevate WNT/β-catenin signaling (i.e. enriched in the Top 10% gate) are shown in ball-and-stick representation. **c**, Structural model of the complex between β-catenin (PDB 2Z6H, sandy brown) and the BCL9 HD2 domain (PDB 2GL7, dark grey)^48^. In the zoomed view, side chains of β-catenin residues targeted by sgRNAs that reduce WNT/β-catenin signaling (i.e. enriched in the Bottom 15% gate) are shown in ball-and-stick representation. **d**, Structural model of the complex between β-catenin (PDB 2Z6H, sandy brown) and the N-terminus of LEF1 (PDB 3OUX, dark grey)^49^. In the zoomed view, side chains of β-catenin residues targeted by sgRNAs that reduce WNT/β-catenin signaling (i.e. enriched in the Bottom 15% gate) are shown in ball-and-stick representation. **e**, Structural model of the complex between β-catenin (PDB 2Z6H, sandy brown) and the N-terminus of TCF4 (1JDH, blue)^50^. Zoomed views are used to show three different regions of this extended interaction interface. Side chains of β-catenin residues targeted by sgRNAs that reduce WNT/β-catenin signaling (i.e. enriched in the Bottom 15% gate) are shown in ball-and-stick representation. Residues numbers circled by a rounded rectangle have been implicated in β-catenin function in prior structural or biochemical studies. Side chains of the partner proteins (β-TRCP, LEF1 or TCF4) are shown if they make a predicted contact with β-catenin. Dashed black lines show putative electrostatic interactions. See also Supplementary Table 2.

**Extended Data Fig. 6.**
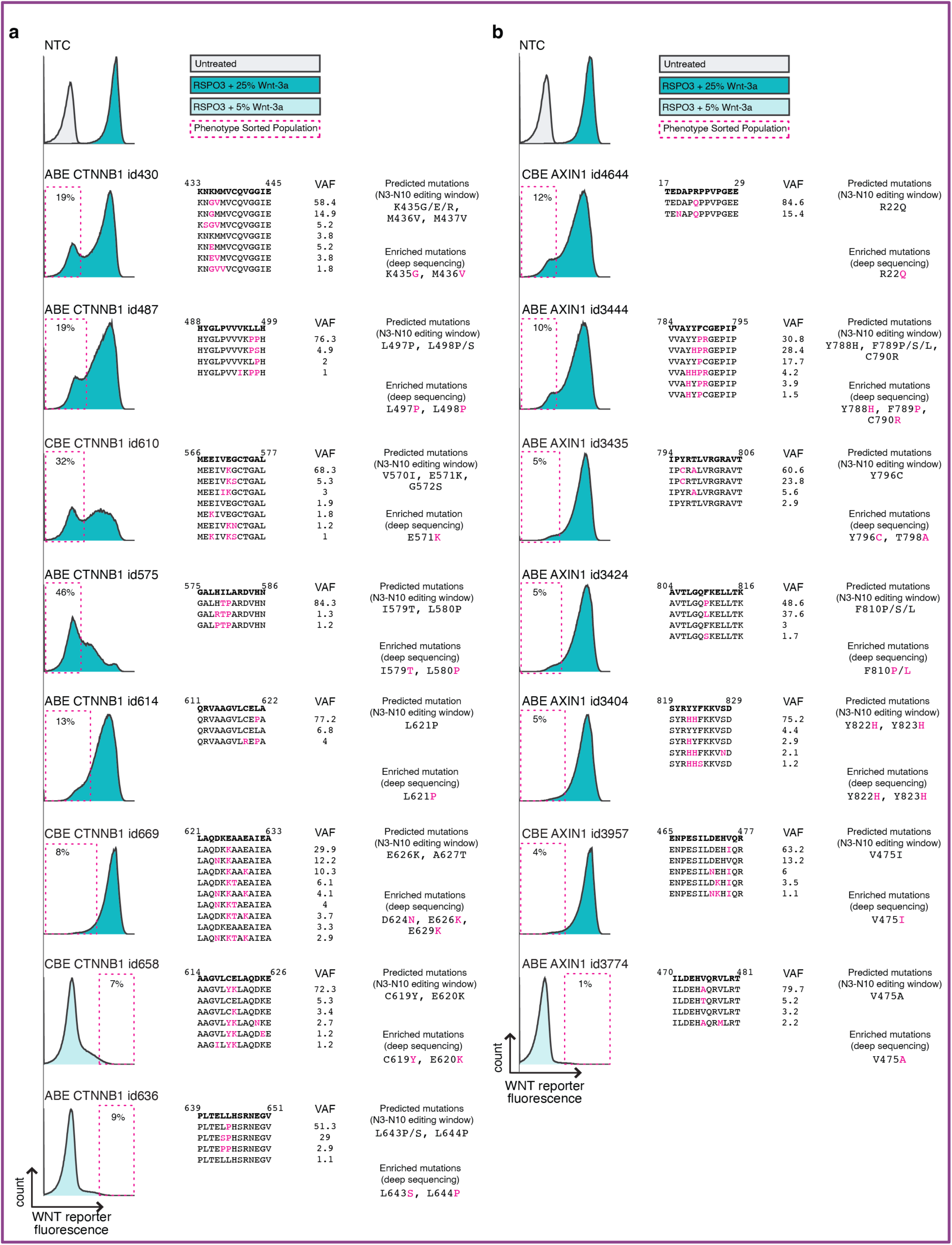
Illumina sequencing strategy to identify sgRNA directed mutations in HEK293T-7TS cells used for mechanistic studies. This figure outlines the strategy for FACS-based enrichment and deep sequencing of base-edited HEK293T-7TS cells. Approximately 1 million cells transfected with pRDA plasmids encoding listed sgRNAs were sorted, expanded, and then subjected to deep sequencing of the targeted locus. **a**, FACS gates (indicated by a dotted magenta rectangle) are shown for cells transfected with sgRNAs targeting *CTNNB1*. Cells were treated with a high dose of ligand (RSPO3 + 25% Wnt-3a conditioned medium) to sort for mutants with reduced WNT reporter fluorescence, or a low dose (RSPO3 + 5% Wnt-3a conditioned medium) to sort for mutants with elevated WNT reporter fluorescence. The right panel shows the predicted mutations in the N3-N10 protospacer window and the measured frequencies of various alleles in the sorted cells (variant allele frequency or VAF). **b**, The same FACS gating strategy is shown for pooled AXIN1 base-edited cell lines. See also Supplementary Table 3.

**Extended Data Fig. 7.**
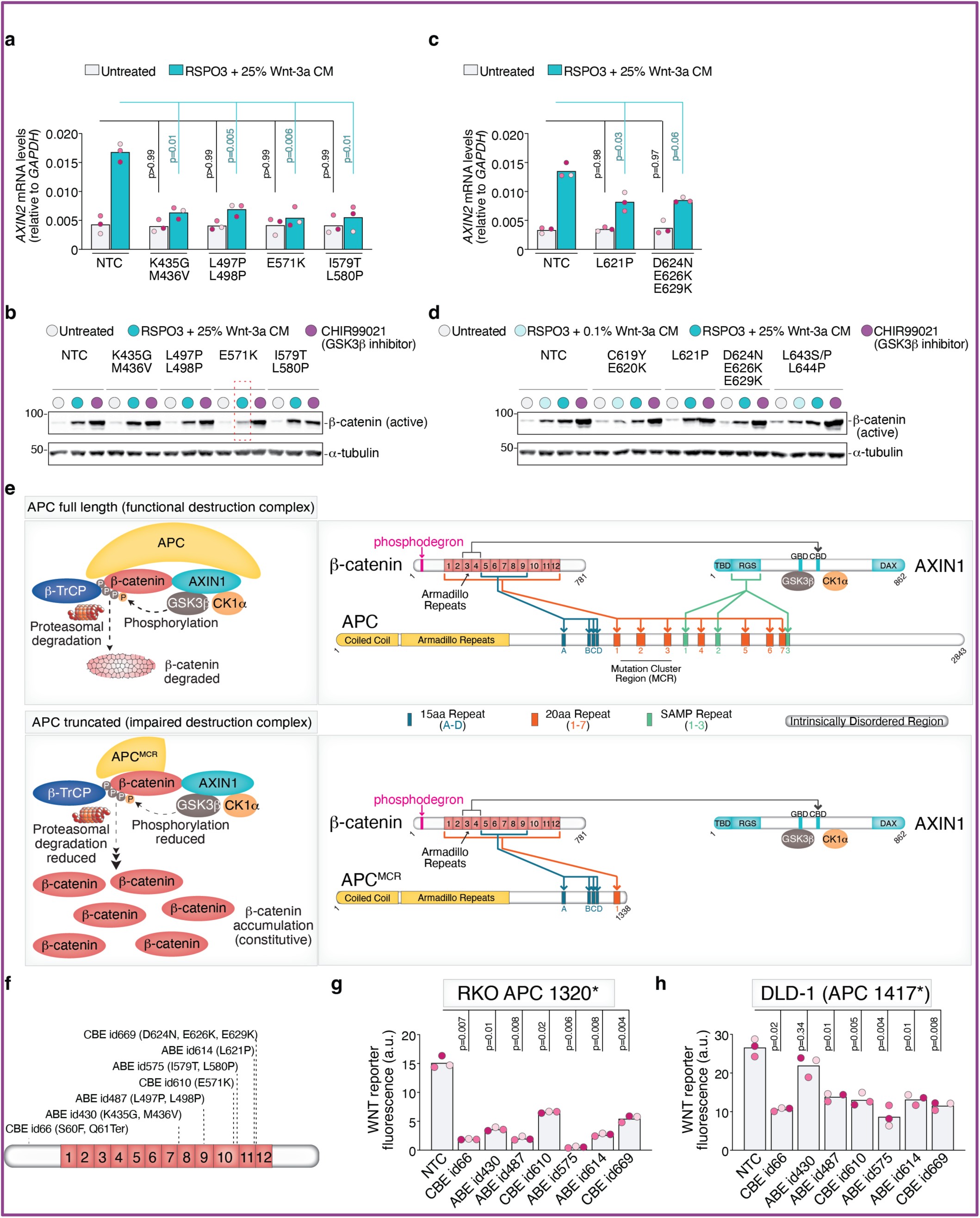
Functional analysis of screen-identified mutations in β-catenin across assays and cell lines. This figure provides supporting data for Fig. 3 and Fig. 4. As an alternative to the synthetic 7TS fluorescent transcriptional reporter (shown in **Fig.1c**), we tested whether sgRNA-induced mutations in β-catenin influence the induction *AXIN2*, an endogenous WNT target gene widely used as a metric for WNT/β-catenin signaling strength, or the abundance of β-catenin. Additionally, we tested whether selected mutations in β-catenin influence the constitutive oncogenic signaling seen in CRC cell lines carrying truncating mutations in APC. **a,c**, *AXIN2* mRNA levels were measured by quantitative reverse-transcription PCR (qRT-PCR) in HEK293T-7TS cells carrying the indicated mutations in β-catenin (confirmed by Illumina sequencing as shown in **Extended Data Fig.6**). Cells were left untreated or treated (24 h) with the indicated concentration of WNTs (RSPO3+Wnt-3a). **b,d,** the abundance of β-catenin protein in these same cell lines was measured by immunoblotting after a 4 h exposure to RSPO3 + Wnt-3a or GSK3β inhibitor (CHIR99021). α-tubulin serves as a loading control. The dashed red rectangle in **b** highlights a β-catenin mutation (E571K) with a defect in Wnt-3a-induced β-catenin accumulation but not CHIR99021-induced accumulation. **e**, PPIs between components of the wild-type, ligand-regulated BDC containing full-length APC (top) and the oncogenic BDC (bottom) containing truncating mutations in APC (APC^mcr^). Colored lines indicate previously mapped PPIs between AXIN1-APC, APC-β-catenin, and AXIN1-β-catenin. Most oncogenic mutations in APC truncate the protein in the Mutation Cluster Region (MCR), eliminating all direct contacts between the APC SAMP motifs and the AXIN1 RGS domain and a subset of contacts between APC and β-catenin. TBD, tankyrase binding domain; DVL, Dishevelled; CBD, β-catenin binding domain of AXIN1; GBD, GSK3β-binding domain, DAX, Domain present in DVL and AXIN. See **Fig.1b** for further details. **f**, Linear cartoon representation of β-catenin showing the 12 ARM repeats and predicted mutations generated by sgRNAs tested in **g** and **h**. **g,h,** Activity of a WNT fluorescent reporter (7xTCF-GFP, same as the 7TS reporter **Fig.1c** but with GFP) in two CRC cell lines expressing the indicated sgRNAs (**f** shows predicted mutations). APC is truncated at amino acid residue 1320 in RKO cells (left) and residue 1417 in DLD1 cells (right), leading to constitutive, ligand-independent WNT/β-catenin signaling and high GFP fluorescence. Bars represent the mean of three biological replicates (*n*=3, each denoted using a different pink shade). Each data point represents the mean fluorescence from approximately 10,000 cells measured by flow cytometry. In **a,c,g,h,** statistical significance was determined using Brown-Forsythe and Welch ANOVA followed by Dunnett’s T3 multiple comparisons test for pairwise comparisons between groups.

**Extended Data Fig. 8.**
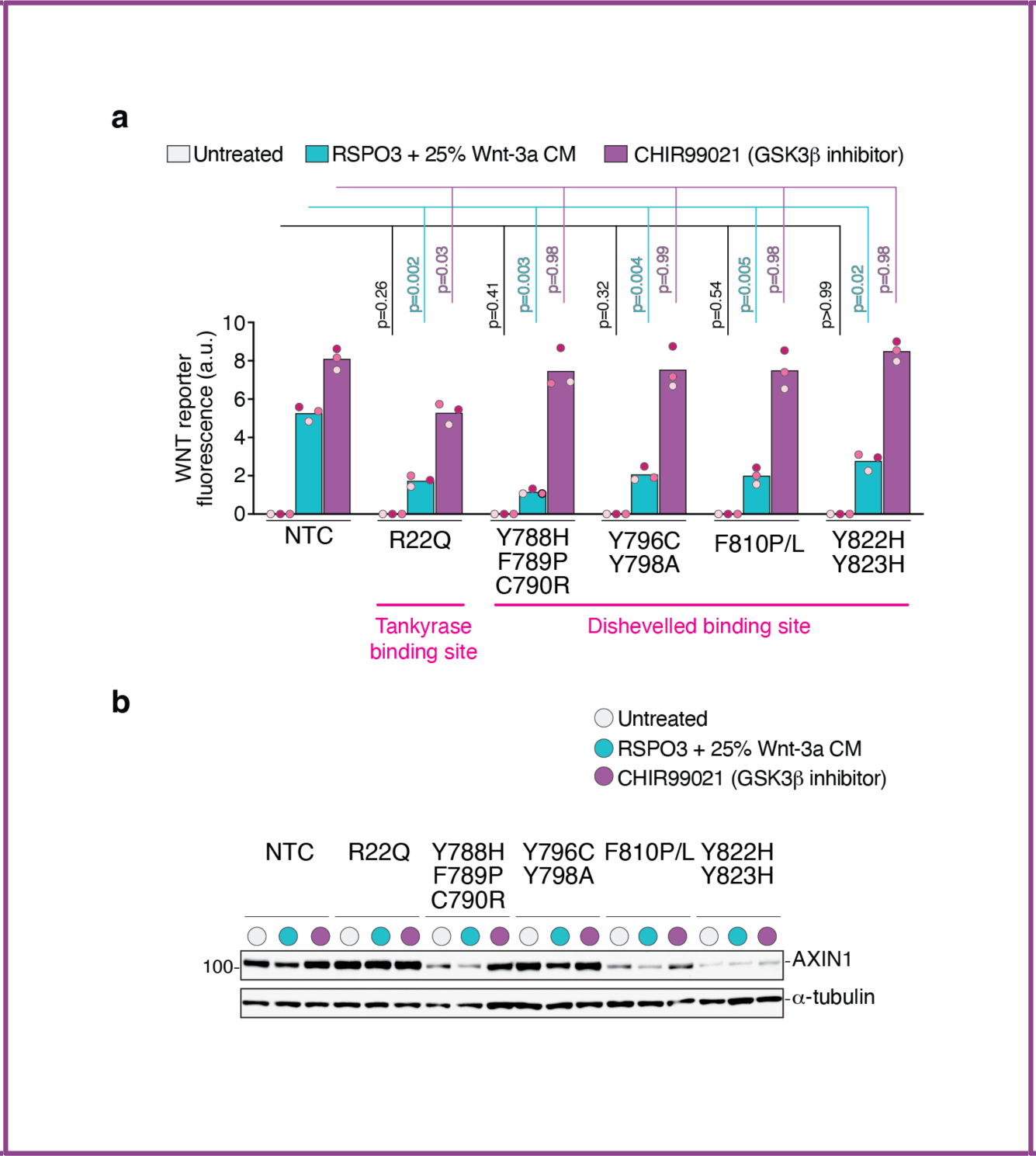
Functional analysis of screen-identified mutations in AXIN1. This figure provides supporting data for Fig.5. **a,** WNT/β-catenin signaling strength was measured in HEK293T-7TS cells carrying the indicated mutations in AXIN1 (confirmed by Illumina sequencing as shown in **Extended Data Fig.6**). Cells were left untreated or treated (24 h) with RSPO3 + Wnt-3a or a GSK3β inhibitor (CHIR99021). Bars represent the mean of three biological replicates, with each replicate indicated by a different pink shade. Each data point represents the mean fluorescence from approximately 10,000 cells. Amino acid residues previously known to be critical for the interaction of AXIN1 with Tankyrase 1 (TNKS1) and amino acid residues predicted to be important for the interaction with the DIX domain of Dishevelled are denoted. These amino acid residues are shown on the structural models in Fig. 5b and 5c. Statistical significance was determined using Brown-Forsythe and Welch ANOVA followed by Dunnett’s T3 multiple comparisons test for pairwise comparisons between groups. **b**, AXIN1 abundance was measured in these same cell lines after cells were left untreated or treated (5 h) with RSPO3 + Wnt-3a or a GSK3β inhibitor (CHIR99021). α-tubulin serves as a loading control.

**Extended Data Fig. 9.**
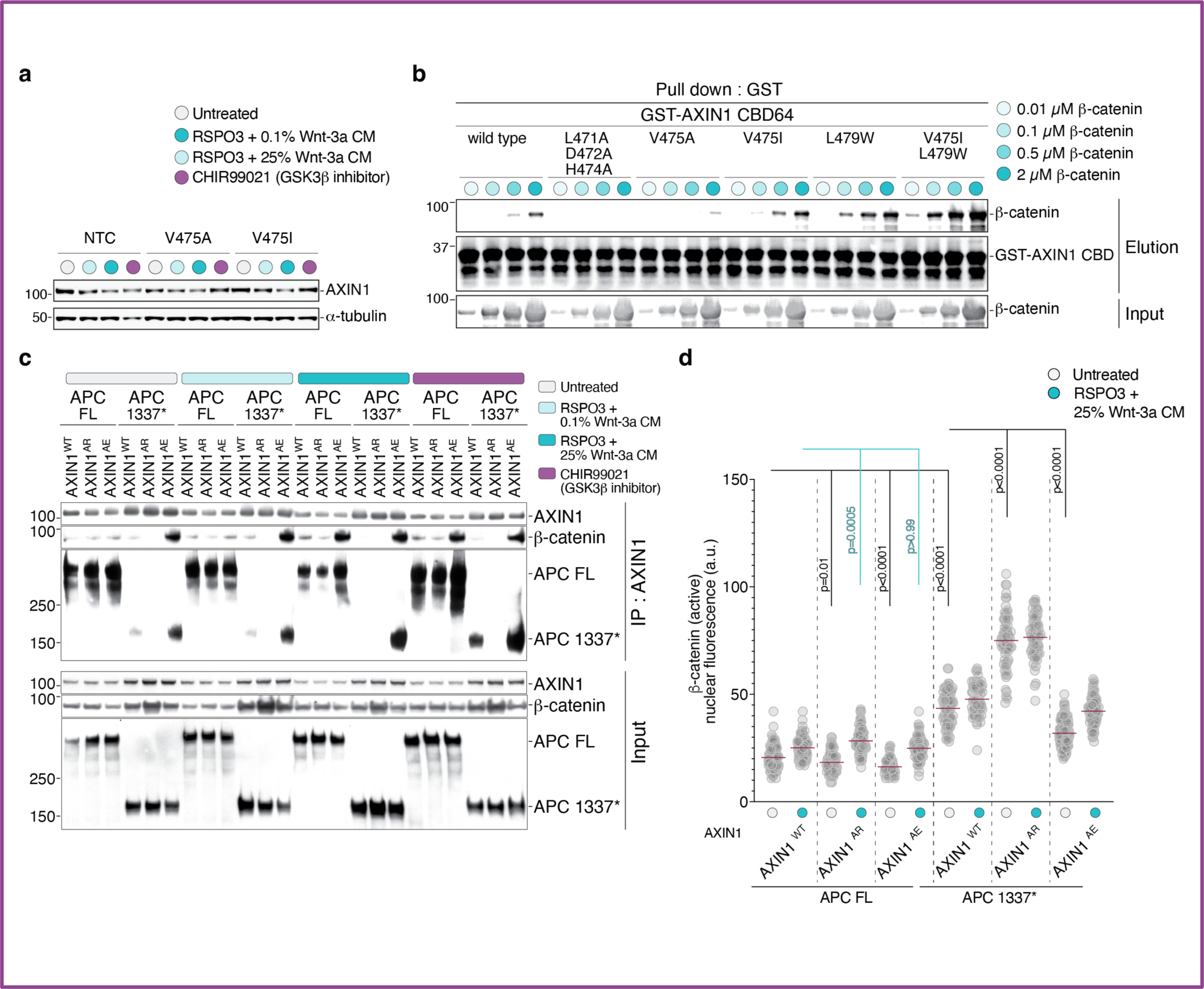
Characterization of AXIN1 mutations in its β-catenin binding domain (CBD). This figure provides supporting data for Fig. 6 and Fig. 7. **a**, AXIN1 protein abundance in HEK293T-7TS cells carrying the indicated mutations (verified by Illumina sequencing) left untreated or treated (5 h) with RSPO3 + Wnt-3a or a GSK3β inhibitor (CHIR99021). α-tubulin serves as a loading control. **b**, An affinity pulldown assay was used as an alternative to Biolayer Interferometry (BLI, **Fig.6d**) to measure the interaction of purified β-catenin with AXIN1 CBD fragments containing the indicated mutations. GST-tagged AXIN1 CBD fragments (1 µM), expressed and purified from *E. coli*, were immobilized on glutathione beads and incubated with increasing concentrations of untagged, full-length, purified β-catenin. β-catenin abundance in the input (5%) and β-catenin captured on the beads (10%) was measured by immunoblotting. **c**, Association of endogenous AXIN1 with APC and β-catenin in extracts from clonal eHAP1 cells measured using an AXIN1 IP followed by immunoblotting. Sequence verified clonal cell lines (**Extended Data Fig.10**) contained the indicated combinations of full-length APC (APC FL) or truncated APC (APC 1337) and one of three AXIN1 variants-- AXIN1^WT^, AXIN1^AE^ (V775I/L479W) and AXIN1^AR^ (L471A/D472A/H474A)-- with varying affinities for β-catenin (see **Fig.6d**). Extracts were made from untreated cells or cells treated (1h) with low-dose Wnt-3a, high-dose Wnt-3a or CHIR99021. This experiment is identical to the one shown in **Fig.7a** but performed using an independent set of clonal cell lines. **d**, Nuclear abundances of active (non-phosphorylated) β-catenin were measured by immunofluorescence and quantitative image analysis in clonal cell lines carrying the indicated combinations of *APC* and *AXIN1* alleles (same cell lines as those used in **Fig.7a** for IP experiments). Measurements were performed in untreated cells or cells treated with RSPO3 + Wnt-3a for 2 h. Statistical significance was determined using Brown-Forsythe and Welch ANOVA followed by Dunnett’s T3 multiple comparisons test for pairwise comparisons between groups.

**Extended Data Fig. 10.**
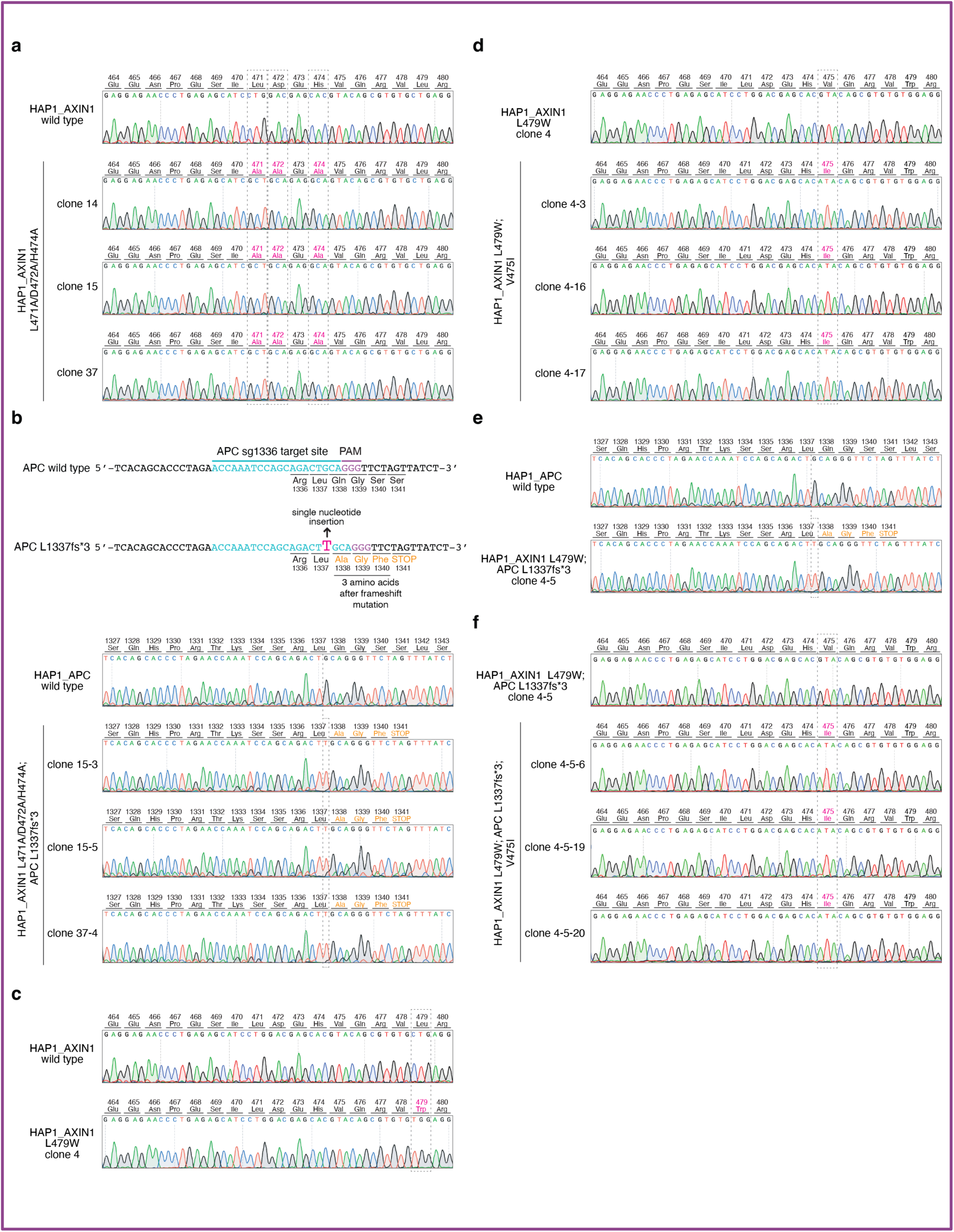
Generation and sequence verification of clonal eHAP1 cell lines carrying various combinations of *AXIN1* and *APC* alleles (used in Fig.7 and Extended Data Fig.9). All mutations were introduced into the single *APC* and *AXIN1* alleles in eHAP1 cells using a combination of CRISPR/Cas9-mediated non-homologous end joining (NHEJ), homology-directed repair (HDR), and base editing. **a**, Sanger sequencing traces confirming the presence mutations (L471A/D472A/H474A) generated using HDR that reduce AXIN1-β-catenin affinity (AXIN1^AR^, **Fig.6c-d**) in three clonal cell lines. **b**, Position of the sgRNA target site and associated PAM sequence in the *APC* gene used to generate an APC truncation mutant (APC 1337*, **Fig.7a**) using NHEJ. Sanger sequence traces from three clonal cell lines show insertion of a single T nucleotide that shifts the translational frame, leading to an in-frame stop codon three amino acids after Leu 1337. **c**, Sanger sequencing trace of a clonal line harboring one of the two mutations (L479W, **Fig.6c-d**) used to generate the affinity-enhancing AXIN1^AE^ variant using HDR. Although only one trace is shown, four additional clonal lines were generated with similar WNT/β-catenin signaling phenotypes. **d**, Sanger sequencing traces of clonal lines harboring both mutations (V475I/L479W) used to generate the affinity-enhancing AXIN1^AE^ variant (**Fig.6c-d**). The V475I mutation was introduced using base-editing of the cell line shown in **c**. **e**, Sanger sequencing traces of a clonal line edited using NHEJ to introduce an APC truncating mutation (APC 1337*) in the background of the *AXIN1* L479W mutation (from **c**). Although only one trace is shown, seven additional clonal lines were generated with similar WNT/β-catenin signaling phenotypes. **f**, Sanger sequencing traces of clonal lines base edited to introduce the V475I mutation in the background of the AXIN1 L479W and APC 1337* mutations (from panel **e**)

## Notes

### Summary of Updates

Supplementary Table 1, Supplementary Table 2, and Supplementary Table 3 are included in this updated version.

